# Listeriolysin O recruits UBA1 to ubiquitinate NLRP3 and suppress inflammasome defence

**DOI:** 10.64898/2026.09.24.754281

**Authors:** Longwei Yuan, An Ding, Yuchao Feng, Xiaoqing Ming, Yuting Hu, Chen Yang, Mengchen Zhang, Yingkang Wang, Zheng Nie, Haibo Wu

**Author notes:** These authors contributed equally to this work (Longwei Yuan, An Ding, Yuchao Feng). Corresponding author E-mail: Haibo Wu; Zheng Nie.

## Abstract

The NLRP3 inflammasome is a central cytosolic defence pathway against infection, yet intracellular pathogens must restrain this response to preserve their replicative niche. Here we identify a ubiquitin-dependent immune-evasion mechanism in which *L. monocytogenes* uses listeriolysin O (LLO) to destabilise NLRP3. The N-terminal PEST-like sequence of LLO acted as an adaptor that recruited the host ubiquitin-activating enzyme UBA1 and promoted K48-linked polyubiquitination of NLRP3. This modification drove proteasomal NLRP3 degradation and limited caspase-1 activation, gasdermin D cleavage, IL-1β/IL-18 release, and macrophage pyroptosis. Deleting the PEST region preserved basal haemolytic activity but impaired NLRP3 ubiquitination, thereby enhancing inflammasome activation during infection in macrophages and *in vivo*. Conversely, UBA1 overexpression suppressed NLRP3-mediated inflammatory responses and supported intracellular bacterial survival. UBA1 depletion restored NLRP3 protein levels, reduced bacterial burden and delayed lethal infection, albeit with increased inflammatory tissue injury. These findings define a ubiquitin-hijacking strategy that operates at the E1 step, whereby a bacterial toxin redirects host ubiquitin activation towards an innate immune sensor.

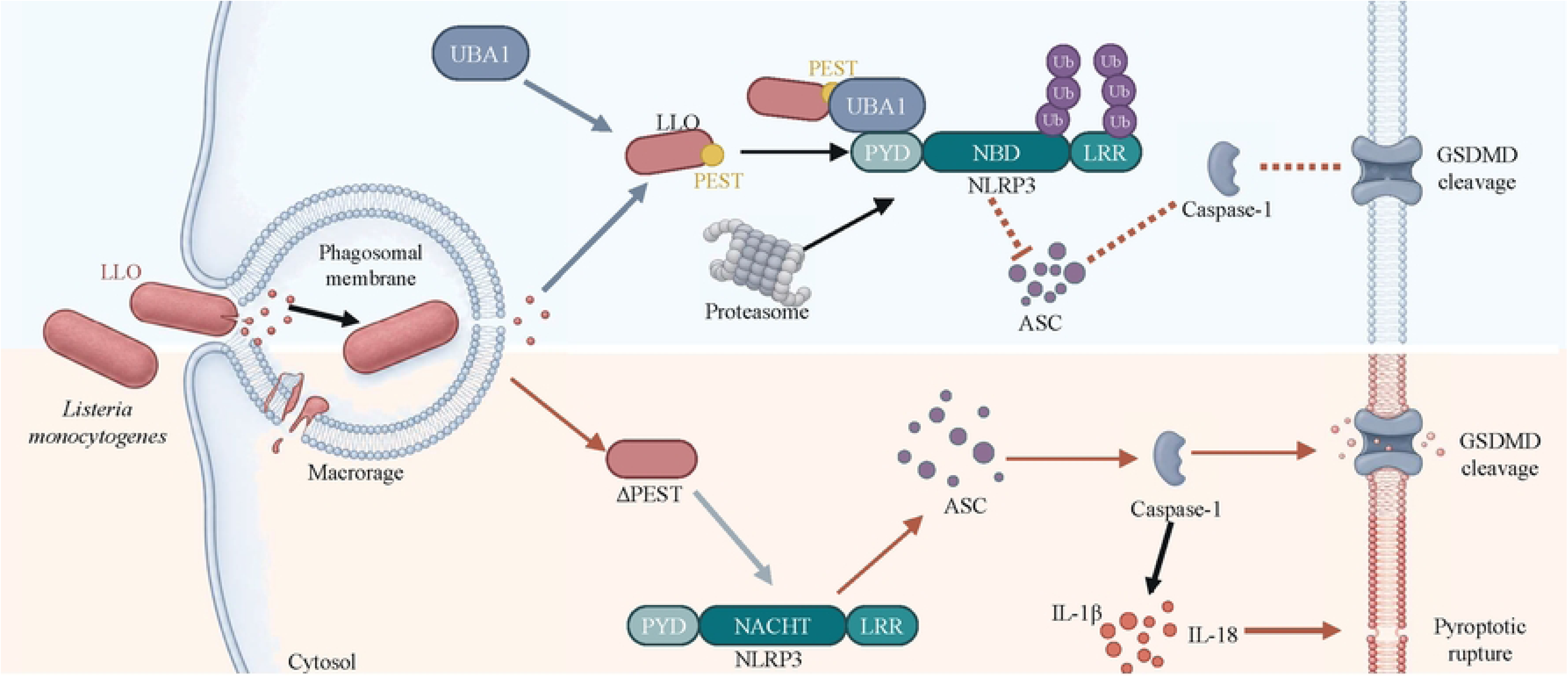

## Introduction

Inflammasomes are cytosolic signalling platforms that convert microbial and damage-associated cues into inflammatory caspase activation. NLRP3 is notable for its ability to sense pore-forming toxins, ionic stress, extracellular ATP and particulate danger signals. Once activated, NLRP3 assembles with ASC and pro-caspase-1 to promote IL-1β and IL-18 maturation^1–5^. Active caspase-1 also cleaves gasdermin D (GSDMD), generating a pore-forming fragment that executes pyroptosis and facilitates cytokine release^6,7^. Although these responses restrict infection, excessive NLRP3 activity contributes to autoinflammatory disease and sterile inflammatory pathology^8–10^. NLRP3 must therefore remain responsive enough to detect infection yet constrained enough to protect the host from inflammatory damage^11,12^.

This balance is governed by transcriptional, conformational and post-translational controls. NF-κB dependent priming licenses NLRP3 expression^13^, whereas ubiquitin editing determines whether NLRP3 is stabilised, assembled or degraded. K48-linked ubiquitination by TRIM31 or SCF-FBXL2 promotes NLRP3 turnover^14,15^. K63-linked ubiquitination mediated by Pellino2 enhances priming-associated responsiveness^16^, and deubiquitination by BRCC3 or ABRO1 permits activation by removing inhibitory ubiquitin marks^17–19^. Additional phosphorylation, palmitoylation and NEK7-dependent structural rearrangements regulate the transition from resting sensor to active inflammasome^20–23^. Together, these studies define NLRP3 as an immune node sensitive to ubiquitin. However, how intracellular bacterial factors exploit this regulatory logic remains incompletely understood.

*L. monocytogenes* (LM) provides a tractable model for this problem because its intracellular lifestyle hinges on a controlled balance between membrane disruption and host-cell preservation. After uptake by phagocytic cells, *L. monocytogenes* escapes the phagosome and replicates in the cytosol, a process requiring the cholesterol-dependent cytolysin LLO encoded by *hly*^24–28^. The N-terminal PEST-like sequence of LLO has been linked to this balance. Deleting this motif preserves pore-forming activity but compromises *L. monocytogenes* pathogenicity, supporting a model in which the PEST region restrains damage caused by the toxin^29^. Subsequent work connected the same sequence to AP-2-dependent endocytic control of plasma-membrane damage during infection^30^. Because PEST-rich sequences in eukaryotic proteins are often associated with ubiquitin-proteasome turnover^31,32^, the LLO PEST region may act as more than a self-limiting toxin element. Whether it can recruit host ubiquitin machinery to remodel an innate immune sensor has remained unexplored.

The ubiquitin cascade begins with the E1 enzyme UBA1, which activates ubiquitin before transfer to E2 enzymes and, through E3 ligases, to substrate proteins, thereby generating ubiquitin signals endowed with substrate and linkage specificity ^33–36^. Many pathogens exploit this pathway, but most characterized bacterial strategies act downstream through E3-ligase mimicry, deubiquitination or redirection of host ubiquitin adaptors^37–43^. The UBA1 inhibitor TAK-243 has underscored the central role of this enzyme in global ubiquitin flux^44^. However, whether pathogens can drive the recruitment of an E1 enzyme to degrade an innate immune sensor remains poorly characterized. Such a mechanism would represent a distinct route by which bacteria suppress host defence upstream of the canonical E2-E3 layer.

Here we show that the PEST region of LLO functions as an adaptor that recruits host UBA1 to promote ubiquitination and degradation of NLRP3. Using PEST-deleted *L. monocytogenes*, reconstitution assays, primary macrophages and systemic infection models, we found that the PEST region was dispensable for basal haemolytic activity but required for NLRP3 ubiquitination via K48 linkages, an event mediated by LLO. LLO assembled a ternary complex with UBA1 and NLRP3 (LLO-UBA1-NLRP3) and promoted proteasomal NLRP3 degradation, thereby dampening pyroptosis and supporting bacterial survival. Together, these findings establish a mechanism whereby a bacterial toxin hijacks ubiquitination at the E1 step to suppress inflammasome-mediated host defence.

## Results

### The LLO PEST domain restrains NLRP3 inflammasome activation

To separate the pore-forming activity of LLO from functions encoded by its PEST domain, we infected primary mouse bone marrow-derived macrophages (BMDMs) with wild-type LM, an LLO-deficient strain (LM_Δ*hly*_) and a strain lacking the LLO PEST domain (LM_ΔPEST_). Mock cells served as controls. This infection panel allowed us to distinguish LLO-dependent cytosolic access from PEST domain-dependent immune regulation.

Deleting the PEST domain did not measurably weaken basal haemolytic activity. Culture supernatants from LM_ΔPEST_ and wild-type LM showed similar sheep red blood cell lysis kinetics and comparable half-haemolytic titres. Recombinant LLO and LLO_ΔPEST_ behaved similarly in parallel assays (S1A-S1C Fig). Thus, differences between wild-type and PEST-deleted bacteria were unlikely to reflect loss of core cytolytic function.

We next examined NLRP3 pathway activation. At 3 h post-infection, LM_Δ*hly*_ induced little NLRP3 accumulation and only weak caspase-1 or GSDMD cleavage. Wild-type LM induced intermediate pathway activation. By contrast, LM_ΔPEST_ markedly increased NLRP3 abundance relative to wild-type LM and enhanced cleavaged caspase-1 and cleavaged GSDMD (Fig 1A). NLRP3 speck quantification showed that specks were rare in mock or LM_Δ*hly*_ -infected BMDMs. Wild-type LM increased speck formation relative to mock infection, whereas LM_ΔPEST_ further increased specks relative to wild-type LM (Fig 1B).

**Fig. 1.**
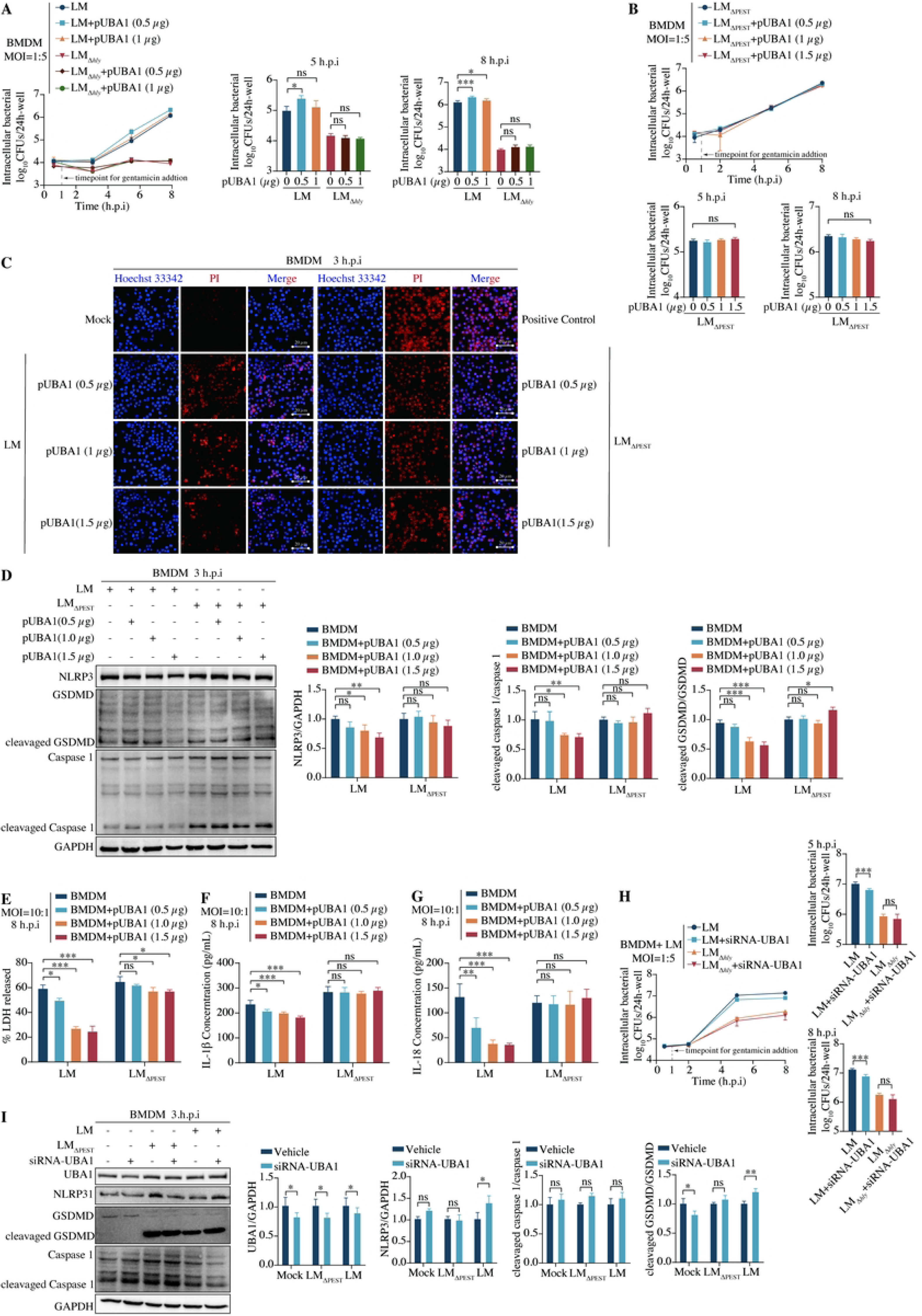
The LLO PEST domain limits NLRP3 inflammasome activation during LM infection. **(A)** Immunoblot analysis of NLRP3 inflammasome activation in BMDMs infected with wild-type LM, LLO-deficient LM (LM_Δ*hly*_) or PEST-deleted LM (LM_ΔPEST_) for 3 h infection. Mock-infected cells served as controls. NLRP3, caspase-1, cleaved caspase-1, GSDMD and cleaved GSDMD were detected, with GAPDH as the loading control. Quantification of NLRP3, cleavaged caspase-1 and cleavaged GSDMD is shown. **(B)** Confocal microscopy analysis of NLRP3 speck formation in infected BMDMs at 3 h.p.i. Cells were stained for NLRP3 (green), intracellular bacteria (magenta) and nuclei (blue). Scale bars, 20 μm. Quantification of NLRP3 speck-positive cells is shown. **(C)** Quantification of LDH release from infected BMDMs at 5 and 8 h.p.i. **(D)** Gentamicin protection assay measuring intracellular bacterial replication in BMDMs infected with different LM mutants. Intracellular bacterial loads were quantified at the 5 and 8 h.p.i.; the infection and gentamicin-treatment timeline is shown. Data are presented as mean ± s.e.m.; ns, not significant; *p < 0.05, **p < 0.01 and ***p < 0.001. In A–D, data are from repeated biological replicates.

Functional readouts were consistent with enhanced inflammasome activation. LDH release was lowest after LM_Δ*hly*_ infection, whereas wild-type LM and LM_ΔPEST_ induced pyroptosis at 5 and 8 h post-infection. The difference between these two LLO-expressing strains increased over time, with LM_ΔPEST_ inducing more LDH release than wild-type LM (Fig 1C). Gentamicin-protection assays showed comparable initial uptake across groups. LM_Δ*hly*_ failed to replicate intracellularly, whereas wild-type LM and LM_ΔPEST_ replicated efficiently, with LM_ΔPEST_ reaching significantly lower burdens than wild-type LM at 5 and 8 h post-infection (Fig 1D). Together, these data show that the LLO PEST domain restrains NLRP3 accumulation, inflammasome assembly and pyroptosis during macrophage infection.

### PEST absence decreases bacterial invasiveness yet aggravates systemic inflammation

We next asked whether the macrophage phenotype was reflected during systemic infection. C57BL/6 mice were injected intravenously with wild-type LM, LM_ΔPEST_ or LM_Δ*hly*_. Tissues were collected 48 h post-infection to measure bacterial burden, inflammatory responses and pathway activation. In addition, a separate cohort was monitored for survival over 14 days (Fig 2A).

**Fig. 2.**
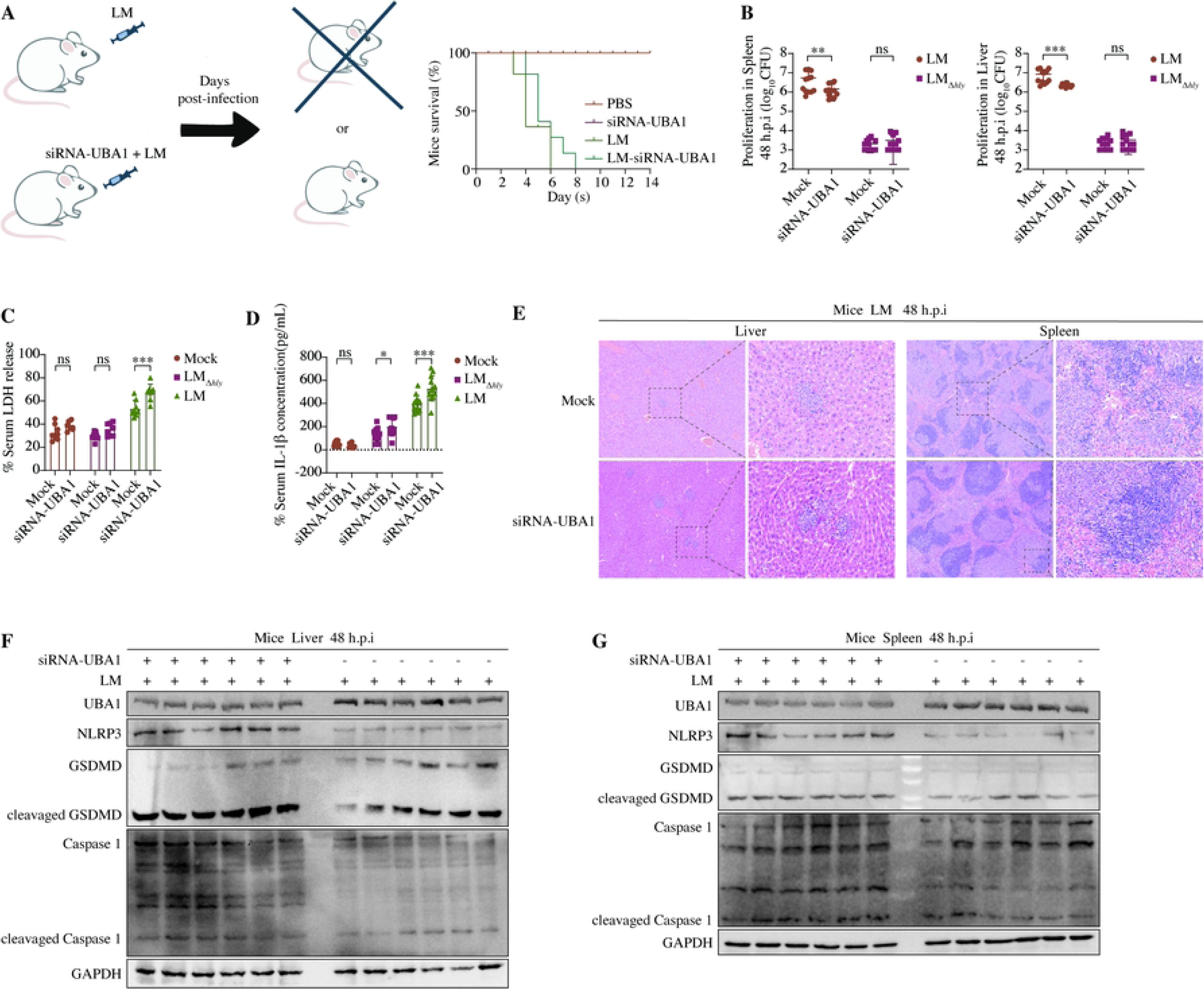
The LLO PEST domain promotes bacterial virulence while limiting systemic inflammasome activation during LM infection. **(A)** Schematic of the systemic *L. monocytogenes* infection model. Mice were infected with LM, LM_ΔPEST_ or LM_Δ*hly*_, and tissues were collected 48 h later for bacterial burden, histology and inflammasome analysis; a separate cohort was monitored for survival. **(B)** Survival of mice infected with the indicated strains over a 14-day period. **(C)** Bacterial burdens in the liver, spleen and brain of infected mice at 48 h.p.i. Bacterial loads are expressed as log10 CFU per tissue. **(D)** Serum LDH levels in infevted mice at 48 h.p.i. **(E)** Serum concentrations of IL-1β and IL-18 in infected mice at 48 h.p.i. **(F)** Representative haematoxylin and eosin staining of liver and spleen sections from mock-infected or infected mice with the indicated bacterial strains at 48 h.p.i. **(G-H)** Immunoblot analysis and densitometric quantification of NLRP3, caspase-1, cleavaged caspase-1, GSDMD and cleavaged GSDMD in the spleen (G) and liver (H) at 48 h.p.i.; GAPDH served as the loading control. Data are presented as mean ± s.e.m.; ns, not significant; *p < 0.05, **p < 0.01 and ***p < 0.001. In B–H, data are from three biological replicates.

As expected, LM_Δ*hly*_ was strongly attenuated and all infected mice survived. Wild-type LM caused rapid death, with all individuals dead by day 7. Compared with wild-type LM, LM_ΔPEST_ delayed mortality, although all mice eventually died by day 9 (Fig 2B). At 48 h post-infection, LM_Δ*hly*_ was barely recovered from the spleen, liver or brain. Despite lacking the PEST motif, LM_ΔPEST_ retained a residual capacity to invade and colonize these organs, although to a significantly diminished extent relative to wild-type LM (Fig 2C). Thus, PEST deletion preserved systemic colonisation capacity but changed the relationship between bacterial burden and acute lethality.

Delayed mortality was accompanied by stronger inflammatory activation. Serum LDH, IL-1β and IL-18 were low in LM_Δ*hly*_ infected mice, whereas LM_ΔPEST_ infection significantly increased these readouts compared with wild-type LM infection (Fig 2D and 2E). Histology supported this pattern: compared with wild-type LM, LM_ΔPEST_ caused more inflammatory-cell recruitment, structural disruption and tissue injury in the liver and spleen (Fig 2F). Immunoblotting further showed that LM_ΔPEST_ infection produced stronger NLRP3 inflammasome activation and more pronounced Caspase 1 and GSDMD cleavage in both organs than wild-type LM (Fig 2G and 2H). Collectively, these findings indicate that the LLO PEST region is critical for LM to establish systemic infection, presumably because the PEST region is required for LM to modulate the NLRP3 inflammasome.

### LLO-PEST drives K48-linked ubiquitination and degradation of NLRP3

Previous studies showed that the N-terminal PEST-like sequence of LLO limits aberrant toxin accumulation in the host cytosol and helps maintain *L. monocytogenes* pathogenicity. The same sequence also promotes AP-2 dependent endocytic clearance of membrane-associated LLO, limiting host-membrane damage^29,30^. In parallel, PEST-rich motifs have long been associated with turnover via the ubiquitin-proteasome system ^31,32^. We therefore asked whether the LLO PEST region influences the stability of the core inflammasome sensor NLRP3, and directly assessed its effects on NLRP3 abundance, proteasome dependent turnover and ubiquitination.

Co-expression of NLRP3 with LLO markedly reduced intracellular NLRP3 abundance, as confirmed by densitometric analysis (Fig 3A). When NLRP3, ubiquitin and LLO were co-expressed and the proteasome was inhibited with MG132, immunoprecipitation showed strong LLO-dependent ubiquitination of NLRP3 (Figs 3B and S2A). These data indicate that LLO promotes proteasomal degradation of NLRP3 in a manner that depends on ubiquitin. Ubiquitin mutants retaining only K6, K11, K27, K29, K33, K48 or K63 were then co-expressed with NLRP3 and LLO. After MG132 treatment, linkage analysis showed that LLO most strongly increased NLRP3 ubiquitination when ubiquitin retained K48, whereas other single-lysine mutants only generated weaker or minimal signals (Figs 3C, 3D, and S2B-S2C). Reciprocal validation showed that K48R ubiquitin abolished the NLRP3 ubiquitination triggered by LLO, whereas K63R had no such effect (Fig 3E).

**Fig. 3.**
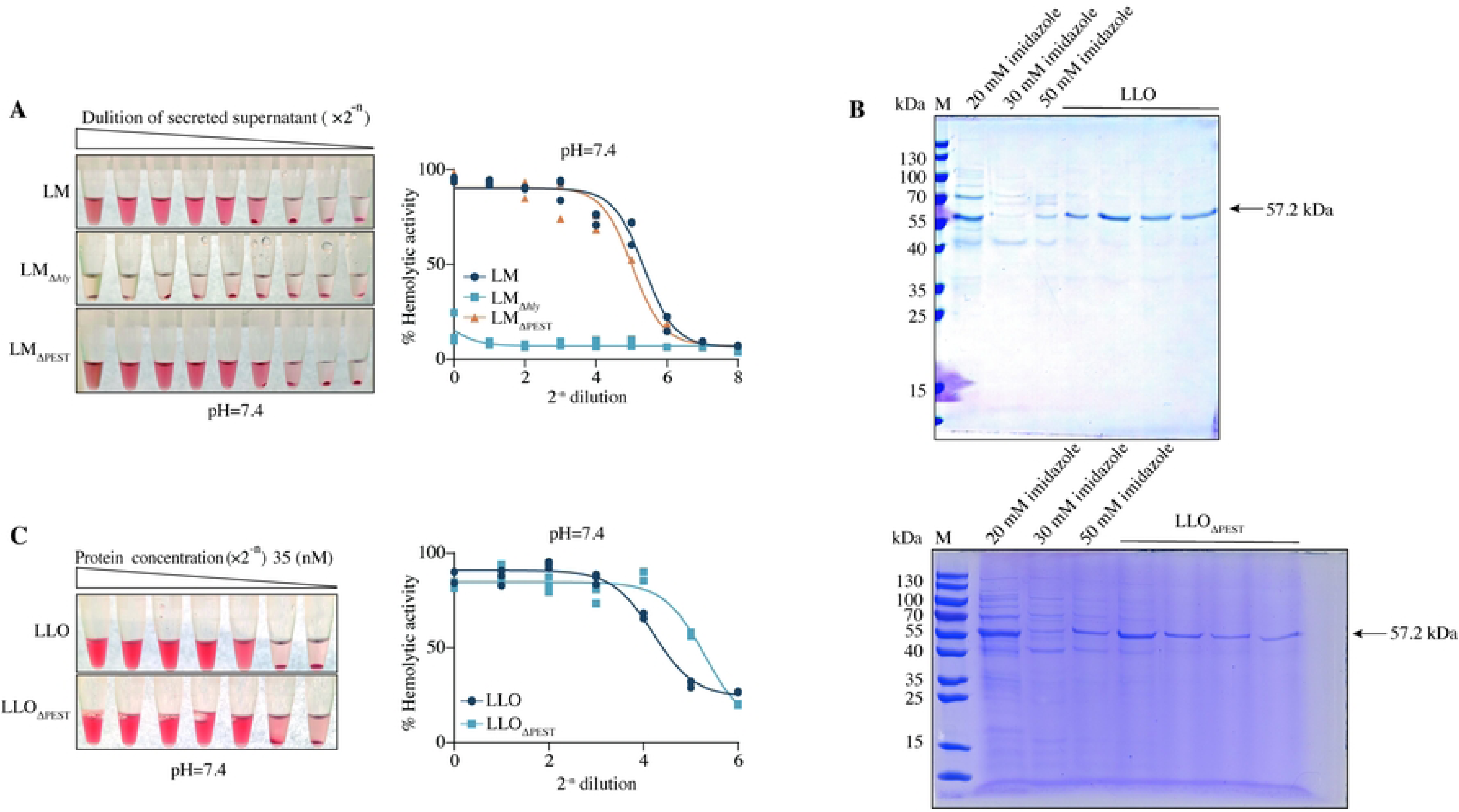
The LLO PEST domain promotes K48-linked ubiquitination and proteasomal degradation of NLRP3. **(A)** Immunoblot analysis of FLAG-NLRP3 abundance in HEK293T cells expressing Myc-LLO, with densitometric quantification of NLRP3 relative to GAPDH. **(B)** Analysis of LLO-induced NLRP3 ubiquitination in MG132-treated HEK293T cells co-expressing FLAG-NLRP3, HA-tagged ubiquitin and Myc-LLO. FLAG-NLRP3 was immunoprecipitated and ubiquitination was detected by HA immunoblotting. **(C)** Identification of ubiquitin linkage specificity induced by LLO. HEK293T cells expressing FLAG-NLRP3 and Myc-LLO were co-transfected with HA-ubiquitin mutants retaining a single lysine residue (K6, K11, K27, K29, K33, K48 or K63), followed by NLRP3 immunoprecipitation and ubiquitination analysis. **(D)** Validation of K48-linked ubiquitination of NLRP3 using wild-type ubiquitin or K48-only ubiquitin mutants in the presence or absence of Myc-LLO. **(E)** Reverse validation using K48- and K63-related ubiquitin mutants to assess the linkage requirement for LLO-induced NLRP3 ubiquitination. **(F)** Comparison of wild-type LLO and LLO_ΔPEST_ in promoting NLRP3 ubiquitination in MG132-treated HEK293T cells. Data are mean ± s.e.m.; **p < 0.01. In A–F, data are from repeated biological replicates.

This activity depended on the PEST domain. After MG132 treatment, wild-type LLO markedly increased NLRP3 ubiquitination, whereas LLO_ΔPEST_ almost completely lost this activity (Figs 3F and S2D). Thus, LLO uses its PEST domain to drive K48-linked polyubiquitination and proteasomal degradation of NLRP3.

### LLO recruits UBA1 to assemble an NLRP3-degrading complex

Given that LLO lacks known catalytic domains or ubiquitin ligase activity, we asked whether it drives NLRP3 degradation by hijacking host ubiquitin-system components. To identify the relevant ubiquitin-system components systematically, we conducted LLO immunoprecipitation followed by mass spectrometry (IP-MS), together with subcellular-localisation annotation, GO/KEGG analysis and degradation-related protein filtering, to generate a set of LLO-associated host candidates (Figs 4A, 4B, and S2E-S2G). In a ranking of ubiquitin system candidates, UBA1, the canonical E1 enzyme at the entry point of the ubiquitin cascade ^33^, emerged as the top-ranked hit. (Fig 4C). Complementary analysis also placed UBA1 within the degradation-related candidate set (S2H Fig). We therefore focused on UBA1 and tested whether it participates in LLO-PEST-dependent NLRP3 ubiquitination and degradation.

**Fig. 4.**
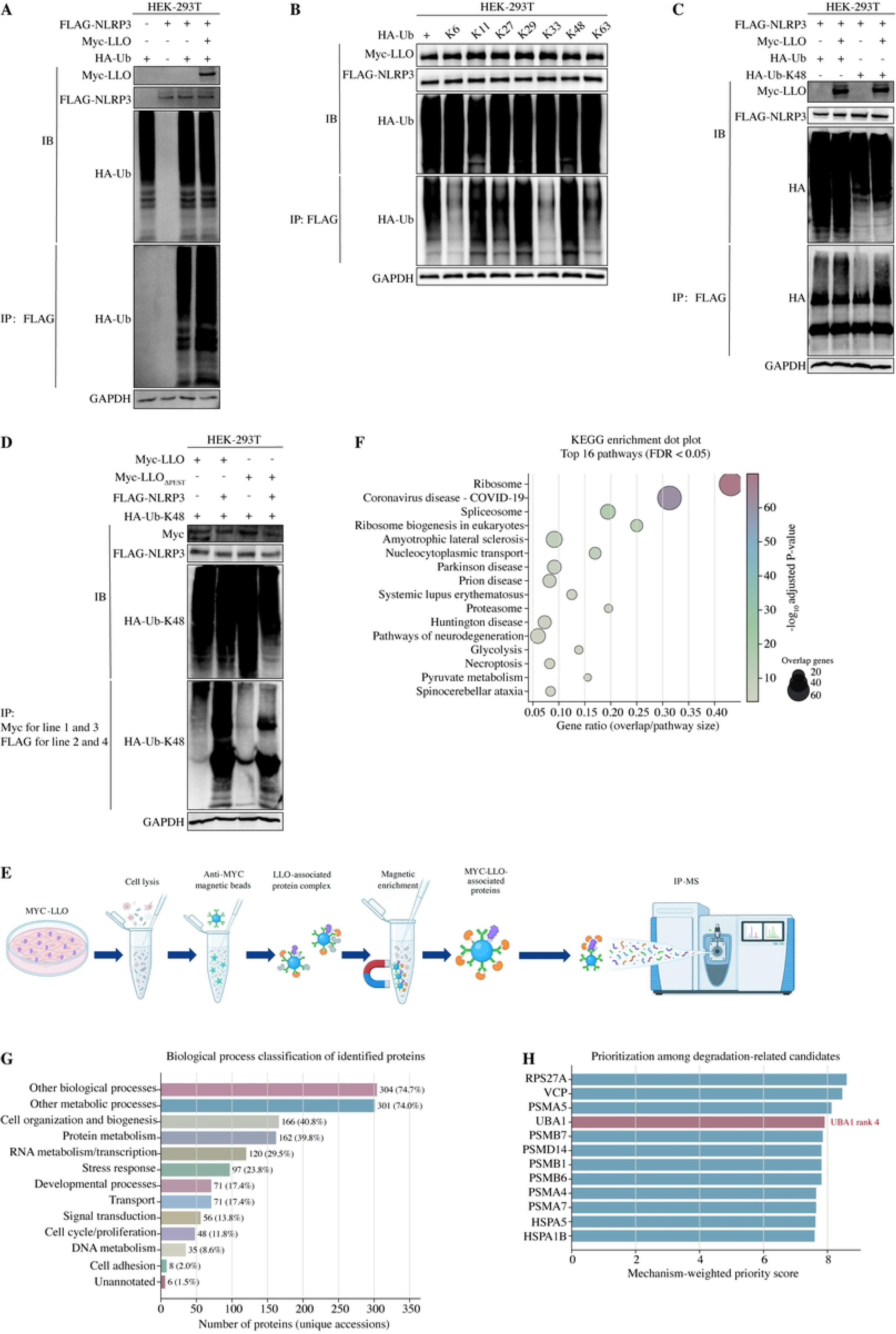
The LLO PEST domain recruits the ubiquitin-activating enzyme UBA1 to facilitate NLRP3 ubiquitination. **(A)** Cellular-component analysis of MYC-LLO-associated proteins identified by affinity purification–mass spectrometry. The circos plot depicts the distribution of total identified proteins and degradation-associated candidates across cellular compartments. **(B)** KEGG pathway enrichment analysis of LLO-associated candidate proteins, showing representative pathways with FDR < 0.05. **(C)** Mechanistically weighted ranking of ubiquitin-system candidates, identifying UBA1 as a prioritised E1 enzyme candidate. **(D-G)** Confocal microscopy analysis of protein co-localization in HEK293T cells. Co-localization of UBA1 with NLRP3 (D) LLO with NLRP3 (E) LLO with UBA1 (F) and triple localization of LLO, UBA1 and NLRP3 (G) were assessed by immunofluorescence staining and line-scan analysis. White lines indicate line-scan paths. Scale bars, 5 μm (E). **(H)** Myc immunoprecipitation analysis showing that Myc-LLO interacts with FLAG-UBA1, whereas Myc-LLO_ΔPEST_ does not bind UBA1 efficiently. **(I)** NLRP3-HA immunoprecipitation analysis demonstrating that LLO promotes the association between NLRP3 and UBA1, whereas this interaction is impaired in the absence of LLO or LLO_ΔPEST_. **(J)** Reciprocal UBA1-FLAG immunoprecipitation validating that wild-type LLO, but not LLO_ΔPEST_, bridges UBA1 and NLRP3. **(K)** Mapping of NLRP3 domains targeted by LLO-induced ubiquitination. LLO preferentially promotes ubiquitination of NLRP3 NBD and LRR regions, whereas LLO_ΔPEST_ produces weaker ubiquitination and degradation signals. **(L)** Domain-specific interaction analysis showing that UBA1 associates with full-length NLRP3 and NLRP3 domain constructs in the presence of LLO, with preferential association with NBD- and LRR-containing regions. **(M)** UBA1 overexpression enhances LLO-mediated NLRP3 ubiquitination/degradation, whereas LLO_ΔPEST_ weakens this effect. **(N)** siRNA-mediated UBA1 knockdown reduces LLO-induced NLRP3 ubiquitination. GAPDH served as a loading control; data are representative of repeated experiments.

To clarify whether UBA1 mediates the PEST-dependent ubiquitination of LLO, we titrated UBA1 overexpression in HEK-293T cells and assessed ubiquitination of LLO relative to LLO_ΔPEST_. Loss of the PEST sequence prevented the increase in LLO ubiquitination normally elicited by UBA1 overexpression (S3A Fig). Confocal imaging showed limited overlap when UBA1 and NLRP3 were expressed alone. By contrast, LLO colocalized with UBA1 and NLRP3 and promoted convergence of the three signals within the same cellular regions (Fig 4D-4G). LLO lacking the PEST region showed markedly reduced colocalization with UBA1 (Fig S3B).

Co-immunoprecipitation validated this candidate prioritization. Wild-type LLO formed a detectable complex with UBA1, whereas LLO_ΔPEST_ failed to bind UBA1 efficiently, indicating that UBA1 recruitment probably depends on the LLO PEST region (Fig 4H). Further showed that NLRP3 co-precipitated with UBA1 only in the presence of wild-type LLO. In the absence of LLO, or when LLO_ΔPEST_ was expressed, NLRP3 failed to pull down UBA1 efficiently (Fig 4I). Reciprocal UBA1 immunoprecipitation produced the same result: UBA1 recovered NLRP3 only in the presence of wild-type LLO, whereas PEST deletion disrupted this interaction (Fig 4J). These findings support a model in which LLO uses its PEST region to bridge UBA1 to NLRP3, rather than UBA1 and NLRP3 spontaneously forming a stable complex.

Domain-level interaction and ubiquitination assays further defined how the complex is organised around NLRP3. NLRP3 domain ubiquitination assays showed that the LRR and NBD regions were the main modules affected by LLO. Wild-type LLO promoted ubiquitination of the NLRP3 LRR and NBD regions and reduced their abundance, whereas LLO_ΔPEST_ generated markedly weaker ubiquitination and degradation signals (Fig 4K). Fragment-interaction assays additionally showed that NLRP3 interacts with residues 26–193 of LLO (Fig.S3C). Domain-mapping co-immunoprecipitation further showed that the NBD and LRR regions of NLRP3 participate in LLO-UBA1-associated complex assembly (Figs 4L and S3D-S3F). Ubiquitination assays probing K48 linkages further supported the conclusion that LLO mainly induces ubiquitination and degradation of the NBD and LRR regions via K48 chains (S3G Fig). Thus, the LLO PEST region most likely behaves as an adaptor, recruiting UBA1 and placing the ubiquitin activation machinery near NLRP3 to drive ubiquitination via K48 chains and proteasomal degradation of NLRP3 modules that contain LRR and NBD.

Functional experiments further established that UBA1 is required for this axis. Under MG132 treatment, wild-type LLO enhanced NLRP3 ubiquitination and reduced NLRP3 protein abundance. UBA1 overexpression further amplified LLO-mediated NLRP3 ubiquitination and degradation, whereas LLO_ΔPEST_ failed to induce this response efficiently (Fig 4M). Conversely, the UBA1 knockdown achieved through siRNA transfection weakened NLRP3 ubiquitination induced by LLO (Fig 4N). These results demonstrate that UBA1 acts upstream of LLO PEST-dependent NLRP3 ubiquitination, but they do not imply that UBA1 itself determines substrate specificity or directly catalyses ubiquitin-chain extension.

To explore possible structural interfaces within this complex, we performed molecular docking of LLO-NLRP3 and LLO-UBA1 pairs. The predicted models identified accessible contact surfaces and candidate interface residues for both pairs (S3H-S3I Fig), complementing our fragment-based mapping, PEST-dependent UBA1 recruitment and co-immunoprecipitation data. Together with the co-immunoprecipitation, these models suggest that LLO may bring UBA1 into sufficient spatial proximity to NLRP3 to support subsequent ubiquitination.. These docking results should be viewed as testable structural hypotheses, not as direct evidence of experimentally resolved interfaces. Taken together, these findings show that LLO exploits its PEST sequence to drive UBA1-dependent ubiquitination and degradation of NLRP3 during LM infection, thereby suppressing NLRP3 inflammasome activation.

### UBA1 mediates PEST-dependent inflammasome suppression in macrophages

We next tested the functional consequences of UBA1 during macrophage infection. Gentamicin-protection assays showed that UBA1 overexpression in BMDMs increased the intracellular burden of wild-type LM at 5 and 8 h post-infection, but there was limited effect on intracellular expansion of LM_Δ*hly*_ (Fig 5A). This pro-survival effect was also confirmed in RAW264.7 macrophages (S4A Fig). By contrast, increasing doses of UBA1 did not promote intracellular growth of LM_ΔPEST_ (Fig 5B). These results indicate that the survival advantage conferred by UBA1 depends on LLO and its PEST region, rather than reflecting a nonspecific effect that promotes growth in all *Listeria* strains.

**Fig. 5.**
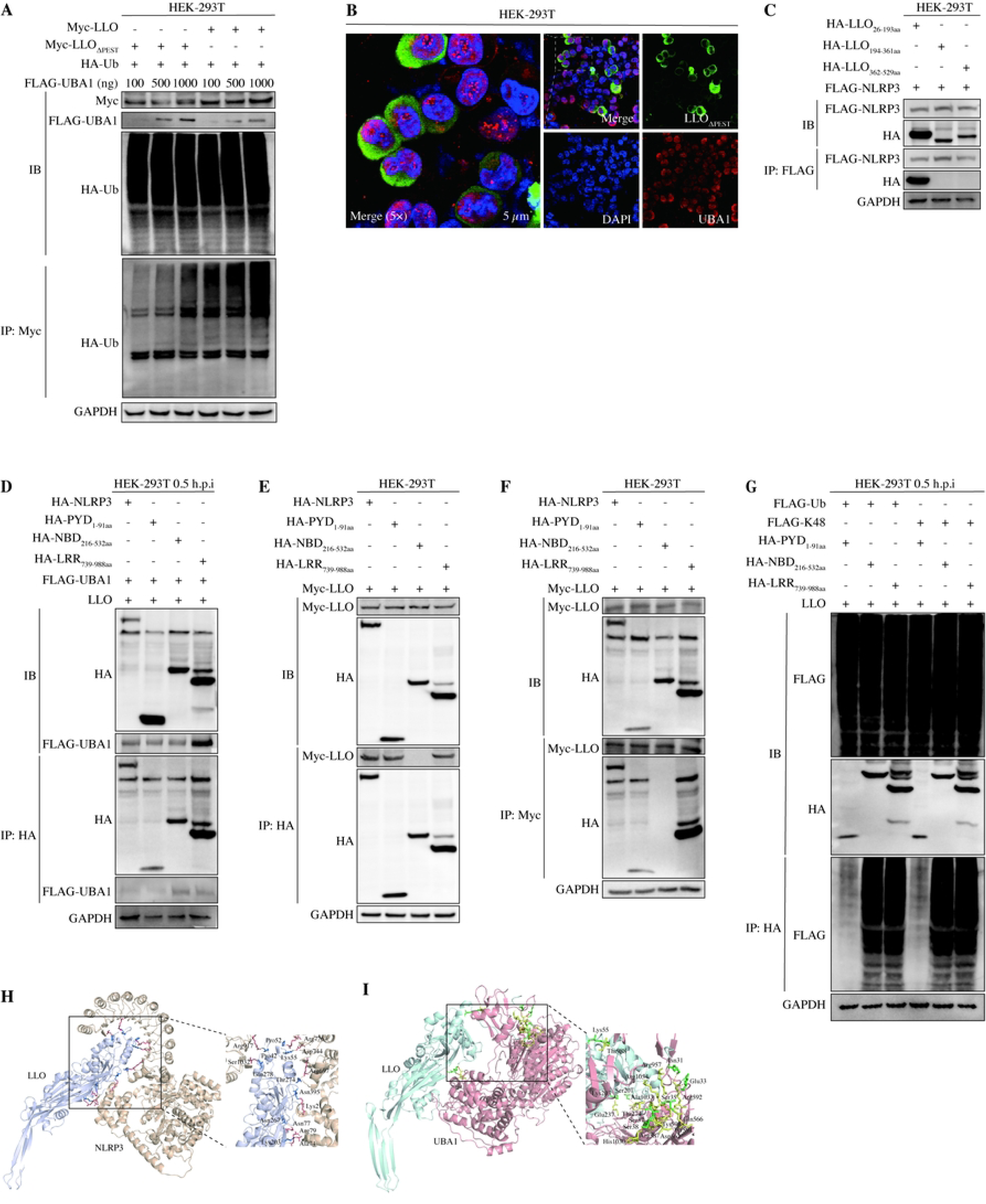
UBA1 mediates LLO PEST domain-dependent NLRP3 degradation and suppression of inflammasome activation in macrophages. **(A)** Gentamicin protection assay measuring intracellular bacterial replication in BMDMs infected with wild-type LM or LM_Δ*hly*_ following UBA1 overexpressing. Increasing amounts of UBA1 plasmid were introduced before infection, and intracellular bacterial loads were quantified at 5 and 8 h.p.i. **(B)** Intracellular bacterial burden of LM_ΔPEST_ in BMDMs overexpressing increasing amounts of UBA1 at 5 and 8 h.p.i. **(C)** Representative Hoechst 33342 and PI staining of BMDMs infected with LM- or LM_ΔPEST_ following UBA1 overexpression. PI-positive cells indicate membrane permeabilization associated with pyroptotic cell death. A positive-control condition is included. Scale bars, 20 μm. **(D)** Immunoblot and quantification showing dose-related effects of UBA1 overexpression on NLRP3 abundance, GSDMD cleavage and caspase-1 cleavage during wild-type LM or LM_ΔPEST_ infection at 3 h.p.i. **(E-G)** Quantification of inflammasome-associated responses in infected BMDMs following UBA1 overexpression. IL-1β secretion (E), IL-18 secretion (F) and LDH release (G) were measured at 8 h.p.i. **(H)** Gentamicin protection assay measuring intracellular bacterial burden of wild-type LM or LM_Δ*hly*_ in BMDMs after siRNA-mediated UBA1 knockdown. **(I)** Immunoblot analysis and densitometric quantification of UBA1, NLRP3, caspase-1, cleavaged caspase-1, GSDMD and cleavaged GSDMD in control and UBA1-depleted BMDMs infected with LM or LM_ΔPEST_. GAPDH served as a loading control. Data are mean ± s.e.m.; ns, not significant; *p < 0.05, **p < 0.01 and ***p < 0.001. In A–I, data are from three biological replicates.

We then examined the effect of UBA1 overexpression on cell death and inflammasome activation. Hoechst 33342/PI staining demonstrated that UBA1 overexpression reduced the proportion of PI-positive cells after wild-type LM infection, whereas cells infected by LM_ΔPEST_ retained strong PI staining (Fig 5C). This result suggests that PEST deletion weakens the ability of UBA1 to limit membrane damage and pyroptosis. Immunoblotting further showed that UBA1 overexpression reduced NLRP3 protein levels and suppressed caspase-1 and GSDMD cleavage during wild-type LM infection. This effect showed a dose-related trend but was markedly weakened or lost during LM_ΔPEST_ infection (Fig 5D). Complementary UBA1 overexpression experiments produced the same pattern: UBA1 suppressed wild-type LM-induced NLRP3 accumulation and cleavage of inflammasome effector proteins, but had limited effects during LM_Δ*hly*_ or LM_ΔPEST_ infection (S4B and S4C Fig).

Downstream inflammatory readouts were consistent with these protein-level changes. LDH-release assays showed that UBA1 overexpression reduced membrane damage induced by wild-type LM, whereas LDH release induced by LM_ΔPEST_ was insensitive to UBA1 overexpression (Fig 5E). UBA1 overexpression reduced mature IL-1β and IL-18 release after wild-type LM infection in a dose-related manner, compared with the modest effect on cytokine release induced by LM_ΔPEST_ (Fig 5F and 5G). ELISA experiments similarly demonstrated that UBA1 mainly suppressed wild-type LM-induced IL-1β and IL-18 secretion, whereas the inhibitory effect was weaker in the LM_Δ*hly*_ and LM_ΔPEST_ groups (S4D and S4E Fig). Notably, IL-1β mRNA increased or fluctuated under wild-type LM with UBA1 conditions, whereas IL-18 mRNA changed little overall (S4F and S4G Fig). UBA1-dependent inhibition of IL-1β and IL-18 release is therefore more consistent with restricted inflammasome-mediated processing and release than with uniform suppression of cytokine transcription.

Finally, we performed siRNA-mediated UBA1 knockdown for reciprocal validation. Immunoblotting across different siRNA sequences and treatment durations confirmed UBA1-knockdown efficiency (S4H Fig). In BMDMs, UBA1 knockdown reduced the intracellular burden of wild-type LM at 5 and 8 h, whereas the effect on LM_Δ*hly*_ was insignificant (Fig 5H), indicating that endogenous UBA1 is required for the LLO-dependent intracellular survival advantage. Consistently, UBA1 knockdown restored NLRP3 levels and enhanced caspase-1/GSDMD cleavage during wild-type LM infection. This effect was weak or absent during LM_ΔPEST_ or LM_Δ*hly*_ infection (Figs 5I and S4I). NLRP3-domain functional experiments further showed that NLRP3 domain integrity affects intracellular LM survival, with LRR deletion causing the most pronounced reduction in bacterial burden (S4J Fig). This pattern is consistent with involvement of the LRR region in LLO-UBA1-associated complex assembly and NLRP3 degradation. Together, these data identify UBA1 as a host factor required for LLO-PEST-dependent NLRP3 degradation, inflammasome suppression and enhanced intracellular bacterial survival.

### UBA1 depletion restores NLRP3 signaling during systemic infection

Having established that UBA1 participates in LLO-PEST-dependent immune suppression in macrophages, we next examined this axis during systemic infection *in vivo*. Mice were treated with siRNA-UBA1 or a control and then infected with *L. monocytogenes*. Survival and infection-related readouts were then monitored (Fig 6A). PBS-treated mice and mice treated with siRNA-UBA1 alone remained alive. Wild-type LM infection caused rapid mortality, whereas UBA1 knockdown delayed death, indicating that UBA1 depletion can slow lethal systemic infection.

**Fig. 6.**
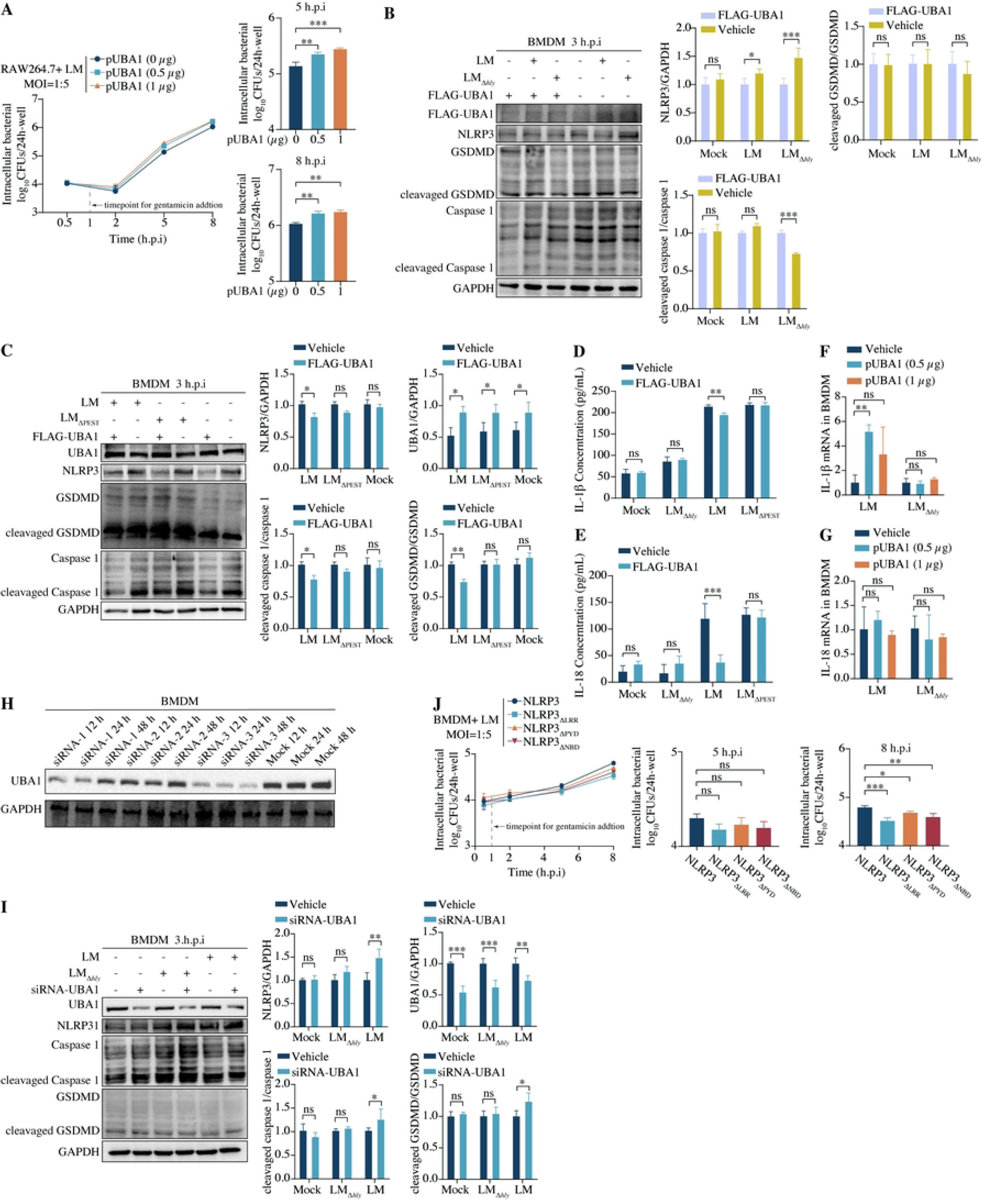
UBA1 depletion restores NLRP3 inflammasome activation and reduces systemic *L. monocytogenes* infection *in vivo*. **(A)** Schematic of *in vivo* UBA1 knockdown and systemic *L. monocytogenes* infection. Mice receiving control or UBA1-targeting siRNA were intravenously infected with wild-type LM, and survival was monitored for 14 days. **(B)** Bacterial burdens in the spleen and liver of control or UBA1-depleted mice infected with wild-type LM or LM_Δ*hly*_ 48h.p.i. Bacterial loads are expressed as log10 colony-forming units (CFU) per organ. **(C-D)** Serum LDH release (C) and IL-1β concentrations (D) measured at 48 h.p.i. **(E)** Representative haematoxylin and eosin staining of the liver and spleen sections from LM-infected mice with or without UBA1 knockdown at 48 h.p.i. Scale bars, 0.1 mm for low-magnification images and 0.02 mm for high-magnification images.**(F-G)** Immunoblot analysis of UBA1, NLRP3, caspase-1, cleavaged caspase-1, GSDMD and cleavaged GSDMD in the liver (F) and spleen (G) from wild-type LM-infected mice with or without UBA1 knockdown at 48 h.p.i.; GAPDH served as the loading control. Data are presented as mean ± s.e.m.; ns, not significant; *p < 0.05, **p < 0.01 and ***p < 0.001.

Consistent with improved host control, UBA1 knockdown reduced wild-type LM bacterial burdens in the spleen and liver, with a more pronounced reduction in the liver. LM_Δ*hly*_ failed to establish effective colonisation regardless of UBA1 knockdown (Fig 6B). At the same time, serum LDH and IL-1β levels increased in UBA1-knockdown mice infected with LM (Fig 6C and 6D), suggesting that restoration of inflammasome responses helps restrict bacterial expansion but is accompanied by stronger cellular injury and inflammatory mediator release.

Histology and immunoblotting further supported this view. After UBA1 knockdown, the liver and spleen from wild-type LM-infected mice showed more pronounced inflammatory infiltration and necrotic lesions (Fig 6E). NLRP3 abundance was restored in infected tissues, and cleavaged caspase-1 and cleavaged GSDMD were increased (Fig 6F and 6G), indicating that UBA1 knockdown *in vivo* relieves LLO-associated suppression of the NLRP3 inflammasome. Control experiments demonstrated that UBA1 knockdown alone did not activate this pathway. LM_Δ*hly*_ infection also failed to induce the NLRP3/GSDMD/caspase-1 activation or tissue pathology observed during wild-type LM infection (S5A-S5C Fig).

Together, the *in vivo* results close the functional loop established our cell-based experiments. *L. monocytogenes* uses UBA1 to connect the LLO PEST region with NLRP3 degradation, thereby suppressing inflammasome activation and promoting infection. Conversely, UBA1 knockdown restores NLRP3 signaling and reduces bacterial burden, although it may also increase inflammatory tissue injury.

## Discussion

Here, we show that LLO recruits UBA1 to promote K48-linked ubiquitination and proteasomal degradation of NLRP3 in a PEST-dependent manner, uncovering a previously unrecognized mechanism of immune evasion. A key finding of this study is that the LLO PEST domain is bifunctional: it restrains toxin-associated membrane damage while also serving as a pathogen-derived adaptor module that couples LLO to the host ubiquitin machinery. Deletion of PEST region preserves basal haemolytic activity but impaires NLRP3 ubiquitination and degradation, resulting in NLRP3 accumulation, caspase-1 activation, gasdermin D cleavage, cytokine release and pyroptosis. Collectively, our data define LLO as a pleiotropic virulence factor that integrates phagosomal escape, intracellular niche maintenance, and evasion of inflammasome-driven immunity^24–30,45–49^.

Previous work has demonstrated that LLO PEST motif is required for full *L. monocytogenes* pathogenicity and helps limit host-cell damage during infection^29,30^. Extending these findings, our data reveal an additional, trans-acting function for this motif: the same region can recruit UBA1 and redirect ubiquitin-dependent turnover towards a host immune sensor—although the full scope of this mechanism warrants further investigation. Our findings establish PEST-dependent UBA1 recruitment, assembly of an LLO-UBA1-NLRP3 complex and consequent NLRP3 degradation. However, the specific residues within the PEST region that mediate binding have not yet been fully delineated, nor has it been determined whether additional host cofactors contribute to the stabilisation of the complex.

Distinct from most virulence factors of pathogens mimicking E3 ligases to orchestrate the ubiquitination pathway, LLO appears to function through recruiting the upstream E1 enzyme UBA1 into proximity with NLRP3. This observation does not necessarily imply that UBA1 alone determines substrate specificity or drives polyubiquitin chain formation. Rather, it points to LLO biasing access to the ubiquitin cascade by positioning ubiquitin-activation machinery near NLRP3, while downstream E2 and E3 components probably remain necessary for substrate selection and K48-linked chain assembly.

This point is particularly relevant to the PEST domain. PEST sequences are commonly viewed as degron-like elements that can be recognised by ubiquitin ligases or their adaptor proteins. An important unresolved question is therefore whether an E3 ligase, or an E3-associated adaptor, is also recruited into the LLO-UBA1-NLRP3 complex. Such a factor could explain how a bacterial toxin bearing PEST sequence couples UBA1 recruitment to selective NLRP3 ubiquitination. Future studies should define the E2-E3 module, test whether it binds LLO-PEST directly or indirectly, and map the NLRP3 lysine residues that accept K48-linked chains.

The apparent discrepancy between *in vitro* macrophage and *in vivo* systemic readouts should be interpreted with caution. In isolated macrophages, deletion of the PEST region reduced the intracellular survival advantage relative to wild-type LM. *In vivo*, however, LM_ΔPEST_ produced higher tissue burdens together with stronger inflammation. Bacterial load in the organs integrates contributions from bacterial replication, dissemination, inflammatory tissue damage, immune-cell recruitment and clearance kinetics. It therefore need not mirror replication within a single macrophage population. Enhanced inflammation and tissue injury induced by LM_ΔPEST_ may alter tissue permissiveness and dissemination in ways that are not captured by the macrophage assay alone.

The *in vivo* data highlight the cost of reversing this immune-evasion mechanism. UBA1 knockdown delayed lethal disease and reduced bacterial burden, consistent with improved host control. However, it also restored LDH release and IL-1β/IL-18 production, as well as inflammatory tissue injury. This dual effect is consistent with an inflammasome pathway that restricts intracellular pathogens while amplifying inflammatory pathology when activation is excessive^1–10,46–49^. Thus, the LLO-PEST-UBA1 axis should be viewed as a bacterial strategy that promotes infection by limiting inflammasome activation. Therapeutic disruption of this pathway would require selective targeting of infection-specific interfaces, rather than broad inhibition of UBA1.

## Methods

### Bacterial strains and culture conditions

All procedures related to pathogenic bacteria were conducted in accordance with Biosafety Level 2 protocols and guidelines. Wild-type *L. monocytogenes* strain EGD-e, the isogenic LLO-deficient mutant (LM_Δ*hly*_) and the LLO PEST-domain deletion mutant (LM_ΔPEST_) were cultured amid Brain Heart Infusion Broth. Bacteria were grown at 37 °C with 5% CO_2_ to mid-log phase, and growth was evaluated by monitoring at an optical density of 600 nm.For macrophage infection, mid-log-phase bacteria were washed and resuspended in phosphate-buffered saline.

### Mice and ethics statement

Animal experimental procedures were approved by by the Laboratory Animal Welfare and Ethics Committee of Chongqing University. All C57BL/6 mice, aged 6-8 weeks, were housed in a specific-pathogen-free (SPF) facility according to standard humane animal husbandry protocols. Each *in vivo* experiment was performed using 50% female and 50% male animals, and results were not expected to be influenced by sex. All mice were maintained under a 12-h light-dark cycle with ad libitum access to regular food and water.

### Cell culture and transfection of eukaryotic expression vecter and siRNA

Bone marrow-derived macrophages were differentiated from mouse bone marrow cells and cultured in the standard basic DMEM containing 20% L929-cell-conditioned medium, 10% fetal bovine serum and antibiotics. HEK293T and RAW264.7 cells were incubated in DMEM supplemented with 10% fetal bovine serum. These cells were grown at 37 °C in a humidified atmosphere con-taining 5% CO_2_. Plasmid and siRNA transfections were performed with commercial lipid-based transfection reagents according to the manufacturers’ instructions. After 24 h of transfection, the cells were collected for further analysis.

### Bacterial infection and gentamicin protection assays

Macrophages were infected with the indicated *L. monocytogenes* strains at a multiplicity of infection of 10. After 1 h, extracellular bacteria were killed with 50ng/mL gentamicin. When the infection time points were reached, cells were lysed and serial dilutions were plated on Brain Heart Infusion Agar with at least 3 independent groups of each time point. Subsequently, all plates were invertedly placed at 37 °C with 5% CO_2_ until the next day for counting bacterial single colonies.

### Western blotting and immunoprecipitation

Cells or tissue homogenates were lysed in detergent-containing buffer supplemented with 1% protease and 1% phosphatase inhibitors. Equal amounts of protein were separated by SDS-PAGE, transferred to PVDF membranes and probed with antibodies against NLRP3, caspase-1, GSDMD, UBA1, epitope tags and loading controls. For immunoprecipitation, clarified lysates were incubated with tag-specific beads or antibodies, then washed to remove unbound proteins and subsequently analysed by immunoblotting.

### Ubiquitination assays

HEK293T cells were co-transfected with FLAG-NLRP3, HA-ubiquitin mutants and Myc-tagged LLO or LLO_ΔPEST_. In several specific experiments,cells were treated with MG132 before being collected as indicated. FLAG-NLRP3 was immunoprecipitated, and ubiquitination was detected by immunoblotting for HA or ubiquitin linkage-specific signals.

### Immunofluorescence microscopy

BMDMs grown on glass coverslips were infected, fixed, permeabilized and stained with antibodies against NLRP3, LLO or UBA1 as indicated. Nuclei were counterstained with DAPI, and images were acquired by confocal microscopy. NLRP3 specks and colocalisation were quantified from images collected with matched acquisition settings.

### Cytotoxicity, cytokine and RNA analyses

LDH release was measured with cell-culture supernatants or serum mixing with the working buffer of the cytotoxicity assay in microplates with at least three independent replicates of each group.After being incubated for 0.5 h with gentle shaking at room temperature, the stop solution was added. Subsequently, absorbance was measured at OD_490 nm_ with a microplate reader. IL-1β and IL-18 concentrations were measured by respective ELISA kits, and eventually absorbance was measured at OD_450 nm_ with a microplate reader.. Total RNA was extracted with TRIzol, reverse transcribed into cDNA and analysed by quantitative PCR. Gene expression was normalised to *Gapdh* using the 2^-ΔΔCt^ method.

### Mouse systemic infection model

For systemic infection, mice were injected intravenously with the indicated *L. monocytogenes* strains. For *in vivo* UBA1 knockdown, mice were administered with siRNA-UBA1 or control siRNA before bacterial injection. Tissues were collected after 48-hour infection for examining bacterial burden, histopathology and immunoblotting. Separate cohorts were monitored for survival.

### Histopathology and haemolysis assays

Tissues were fixed, paraffin embedded, sectioned and stained with haematoxylin and eosin. For haemolysis assays, bacterial culture supernatants or recombinant LLO proteins were incubated with sheep erythrocytes, and haemoglobin release was measured by absorbance at OD_540 nm_.

### Structure analysis

Structural information for the indicated proteins was obtained from the Protein Data Bank (https://www.rcsb.org/). Molecular docking simulations were performed using DMFold, and structural figures were generated with PyMOL.

### Statistical analysis

Statistical analysis was performed using GraphPad Prism Software. For experiments using cell lines, the data represent at least three independent biological replicates (n = 3). For experiments involving ani-mals, all data were obtained from at least two independent experimental repetitions, with a minimum of six animals per group (n ≥ 6). Data are presented as mean ± s.d. or mean ± s.e.m. as indicated in the figure legends. Two-group comparisons were performed using two-tailed unpaired Student’s t test. One-way analysis of variance (ANOVA) with appropriate multiple comparisons tests was used to compare three independent groups. Survival curves were analysed by the log-rank test. *P* < 0.05 was considered statistically significant. Specific details of statistical analysis are presented above or in associated figure legends.

### Key resources

Supplier and catalogue information was cross-checked against the laboratory procurement list. Where several candidate products were present, all matching records are listed and the product used in the reported experiment must be confirmed from experimental logs. Working dilutions, construct identifiers and other details not captured by the procurement list remain to be completed before submission.

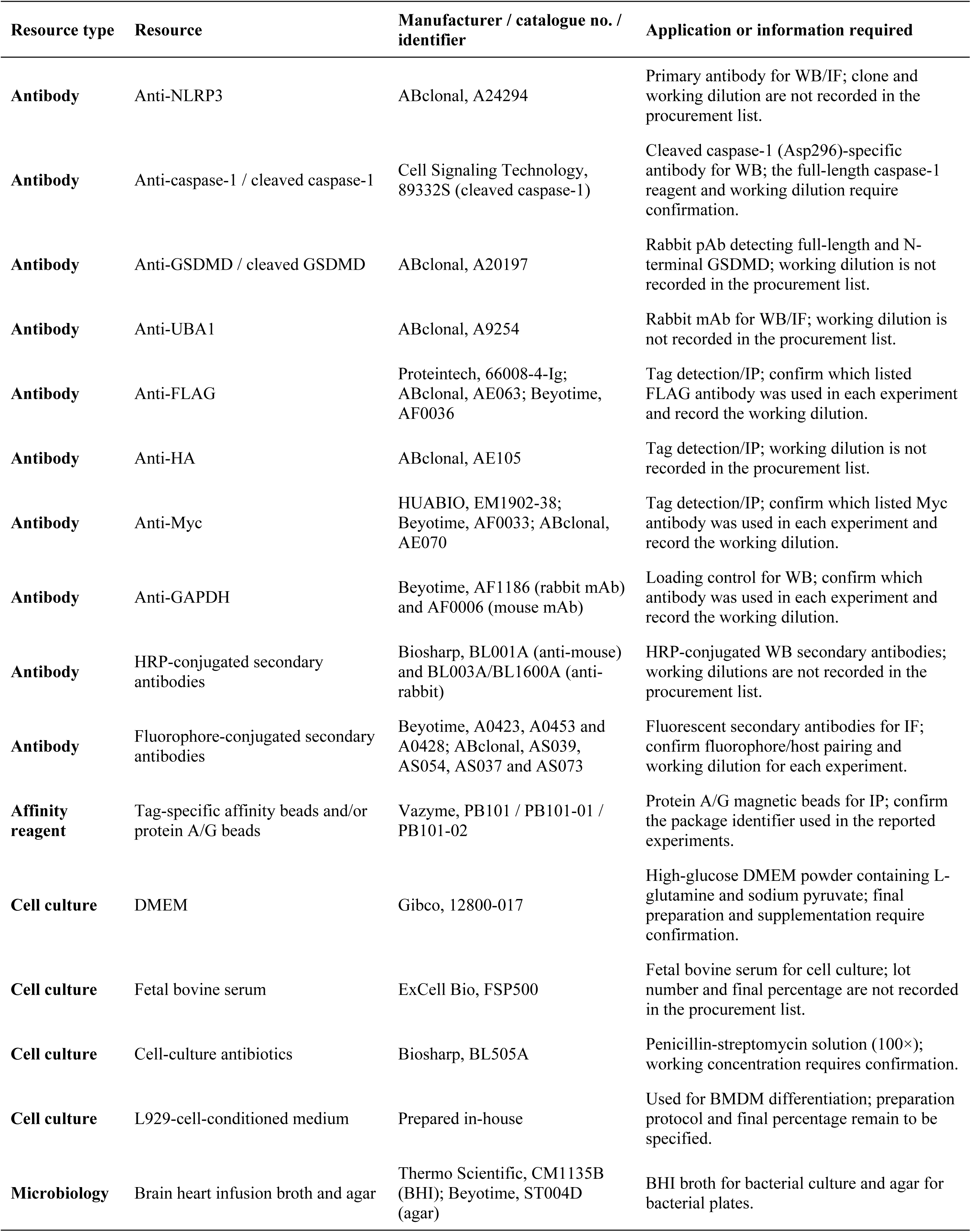
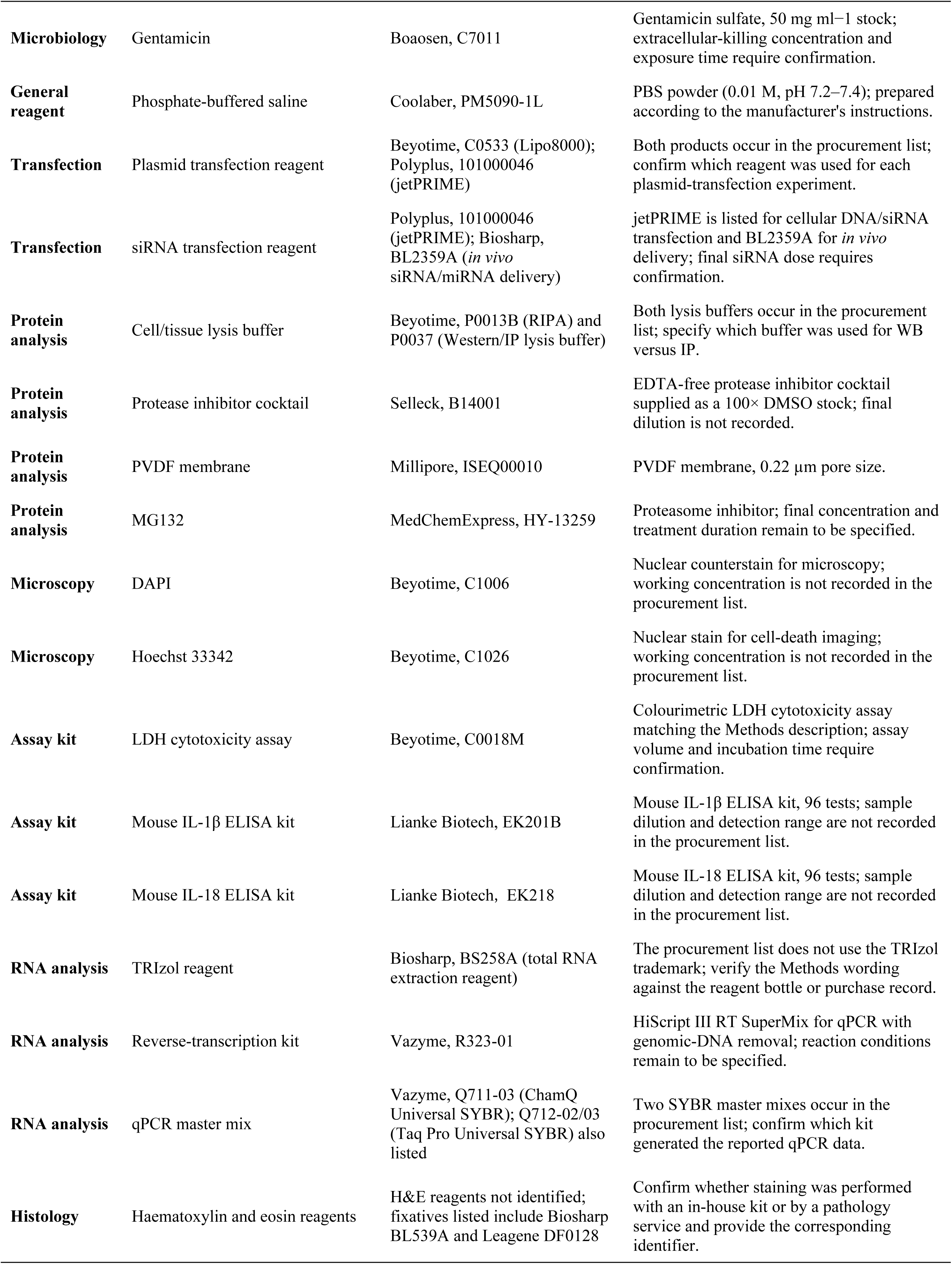

| Resource type | Resource | Manufacturer / catalogue no. / identifier | Application or information required |
| --- | --- | --- | --- |
| Antibody | Anti-NLRP3 | ABclonal, A24294 | Primary antibody for WB/IF; clone and working dilution are not recorded in the procurement list. |
| Antibody | Anti-caspase-1 / cleaved caspase-1 | Cell Signaling Technology, 89332S (cleaved caspase-1) | Cleaved caspase-1 (Asp296)-specific antibody for WB; the full-length caspase-1 reagent and working dilution require confirmation. |
| Antibody | Anti-GSDMD / cleaved GSDMD | ABclonal, A20197 | Rabbit pAb detecting full-length and N-terminal GSDMD; working dilution is not recorded in the procurement list. |
| Antibody | Anti-UBA1 | ABclonal, A9254 | Rabbit mAb for WB/IF; working dilution is not recorded in the procurement list. |
| Antibody | Anti-FLAG | Proteintech, 66008-4-Ig; ABclonal, AE063; Beyotime, AF0036 | Tag detection/IP; confirm which listed FLAG antibody was used in each experiment and record the working dilution. |
| Antibody | Anti-HA | ABclonal, AE105 | Tag detection/IP; working dilution is not recorded in the procurement list. |
| Antibody | Anti-Myc | HUABIO, EM1902-38; Beyotime, AF0033; ABclonal, AE070 | Tag detection/IP; confirm which listed Myc antibody was used in each experiment and record the working dilution. |
| Antibody | Anti-GAPDH | Beyotime, AF1186 (rabbit mAb) and AF0006 (mouse mAb) | Loading control for WB; confirm which antibody was used in each experiment and record the working dilution. |
| Antibody | HRP-conjugated secondary antibodies | Biosharp, BL001A (anti-mouse) and BL003A/BL1600A (anti-rabbit) | HRP-conjugated WB secondary antibodies; working dilutions are not recorded in the procurement list. |
| Antibody | Fluorophore-conjugated secondary antibodies | Beyotime, A0423, A0453 and A0428; ABclonal, AS039, AS054, AS037 and AS073 | Fluorescent secondary antibodies for IF; confirm fluorophore/host pairing and working dilution for each experiment. |
| Affinity reagent | Tag-specific affinity beads and/or protein A/G beads | Vazyme, PB101 / PB101-01 / PB101-02 | Protein A/G magnetic beads for IP; confirm the package identifier used in the reported experiments. |
| Cell culture | DMEM | Gibco, 12800-017 | High-glucose DMEM powder containing L-glutamine and sodium pyruvate; final preparation and supplementation require confirmation. |
| Cell culture | Fetal bovine serum | ExCell Bio, FSP500 | Fetal bovine serum for cell culture; lot number and final percentage are not recorded in the procurement list. |
| Cell culture | Cell-culture antibiotics | Biosharp, BL505A | Penicillin-streptomycin solution (100×); working concentration requires confirmation. |
| Cell culture | L929-cell-conditioned medium | Prepared in-house | Used for BMDM differentiation; preparation protocol and final percentage remain to be specified. |
| Microbiology | Brain heart infusion broth and agar | Thermo Scientific, CM1135B (BHI); Beyotime, ST004D (agar) | BHI broth for bacterial culture and agar for bacterial plates. |
| Microbiology | Gentamicin | Boaosen, C7011 | Gentamicin sulfate, 50 mg ml<sup>-1</sup> stock; extracellular-killing concentration and exposure time require confirmation. |
| General reagent | Phosphate-buffered saline | Coolaber, PM5090-1L | PBS powder (0.01 M, pH 7.2–7.4); prepared according to the manufacturer's instructions. |
| Transfection | Plasmid transfection reagent | Beyotime, C0533 (Lipo8000); Polyplus, 101000046 (jetPRIME) | Both products occur in the procurement list; confirm which reagent was used for each plasmid-transfection experiment. |
| Transfection | siRNA transfection reagent | Polyplus, 101000046 (jetPRIME); Biosharp, BL2359A (*in vivo* siRNA/miRNA delivery) | jetPRIME is listed for cellular DNA/siRNA transfection and BL2359A for *in vivo* delivery; final siRNA dose requires confirmation. |
| Protein analysis | Cell/tissue lysis buffer | Beyotime, P0013B (RIPA) and P0037 (Western/IP lysis buffer) | Both lysis buffers occur in the procurement list; specify which buffer was used for WB versus IP. |
| Protein analysis | Protease inhibitor cocktail | Selleck, B14001 | EDTA-free protease inhibitor cocktail supplied as a 100× DMSO stock; final dilution is not recorded. |
| Protein analysis | PVDF membrane | Millipore, ISEQ00010 | PVDF membrane, 0.22 µm pore size. |
| Protein analysis | MG132 | MedChemExpress, HY-13259 | Proteasome inhibitor; final concentration and treatment duration remain to be specified. |
| Microscopy | DAPI | Beyotime, C1006 | Nuclear counterstain for microscopy; working concentration is not recorded in the procurement list. |
| Microscopy | Hoechst 33342 | Beyotime, C1026 | Nuclear stain for cell-death imaging; working concentration is not recorded in the procurement list. |
| Assay kit | LDH cytotoxicity assay | Beyotime, C0018M | Colourimetric LDH cytotoxicity assay matching the Methods description; assay volume and incubation time require confirmation. |
| Assay kit | Mouse IL-1β ELISA kit | Lianke Biotech, EK201B | Mouse IL-1β ELISA kit, 96 tests; sample dilution and detection range are not recorded in the procurement list. |
| Assay kit | Mouse IL-18 ELISA kit | Lianke Biotech, EK218 | Mouse IL-18 ELISA kit, 96 tests; sample dilution and detection range are not recorded in the procurement list. |
| RNA analysis | TRIzol reagent | Biosharp, BS258A (total RNA extraction reagent) | The procurement list does not use the TRIzol trademark; verify the Methods wording against the reagent bottle or purchase record. |
| RNA analysis | Reverse-transcription kit | Vazyme, R323-01 | HiScript III RT SuperMix for qPCR with genomic-DNA removal; reaction conditions remain to be specified. |
| RNA analysis | qPCR master mix | Vazyme, Q711-03 (ChamQ Universal SYBR); Q712-02/03 (Taq Pro Universal SYBR) also listed | Two SYBR master mixes occur in the procurement list; confirm which kit generated the reported qPCR data. |
| Histology | Haematoxylin and eosin reagents | H&E reagents not identified; fixatives listed include Biosharp BL539A and Leagene DF0128 | Confirm whether staining was performed with an in-house kit or by a pathology service and provide the corresponding identifier. |

## Acknowledgements

This work was supported by the National Natural Science Foundation of China (No.92369115, 82422048, 92469110, and 824B2067), the National Key Research and Development Program of China (No. 2024YFC2310805 and 2021YFC2301405), the Fundamental Research Funds for the Central Universities (2025CDJ-IAISYB-017), and Chongqing Talents: Exceptional Young Talents Project (No. cstc2021ycjh-bgzxm0099). The funders had no role in study design, data collection and analysis, decision to publish, or preparation of the manuscript.

## Figure legends

**Supplementary Fig. 1.**
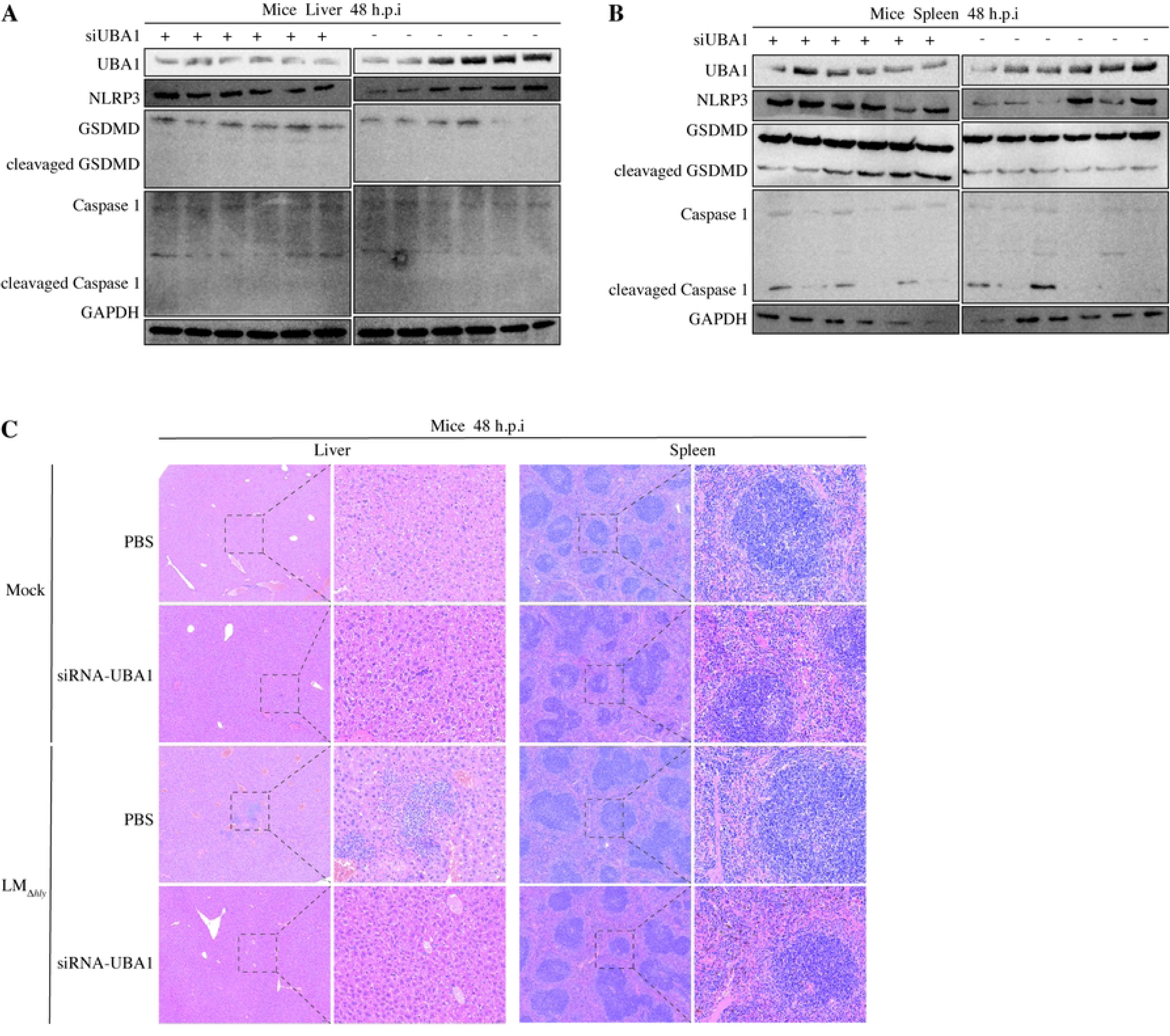
PEST deletion does not abolish basal LLO haemolytic activity. **(A)** Haemolytic activity of culture supernatants from wild-type LM, LM_Δ*hly*_ and LM_ΔPEST_ measured across serial dilutions at pH 7.4. Representative images and quantification of haemolytic activity are shown. **(B)**, SDS-PAGE analysis of purified recombinant LLO and LLO_ΔPEST_ proteins. Representative fractions eluted with the indicated imidazole concentrations are shown. **(C)** Haemolytic activity of recombinant LLO and LLO_ΔPEST_ proteins measured across serial dilutions at pH 7.4. Representative images and quantification of haemolytic activity are shown.

**Supplementary Fig. 2.**
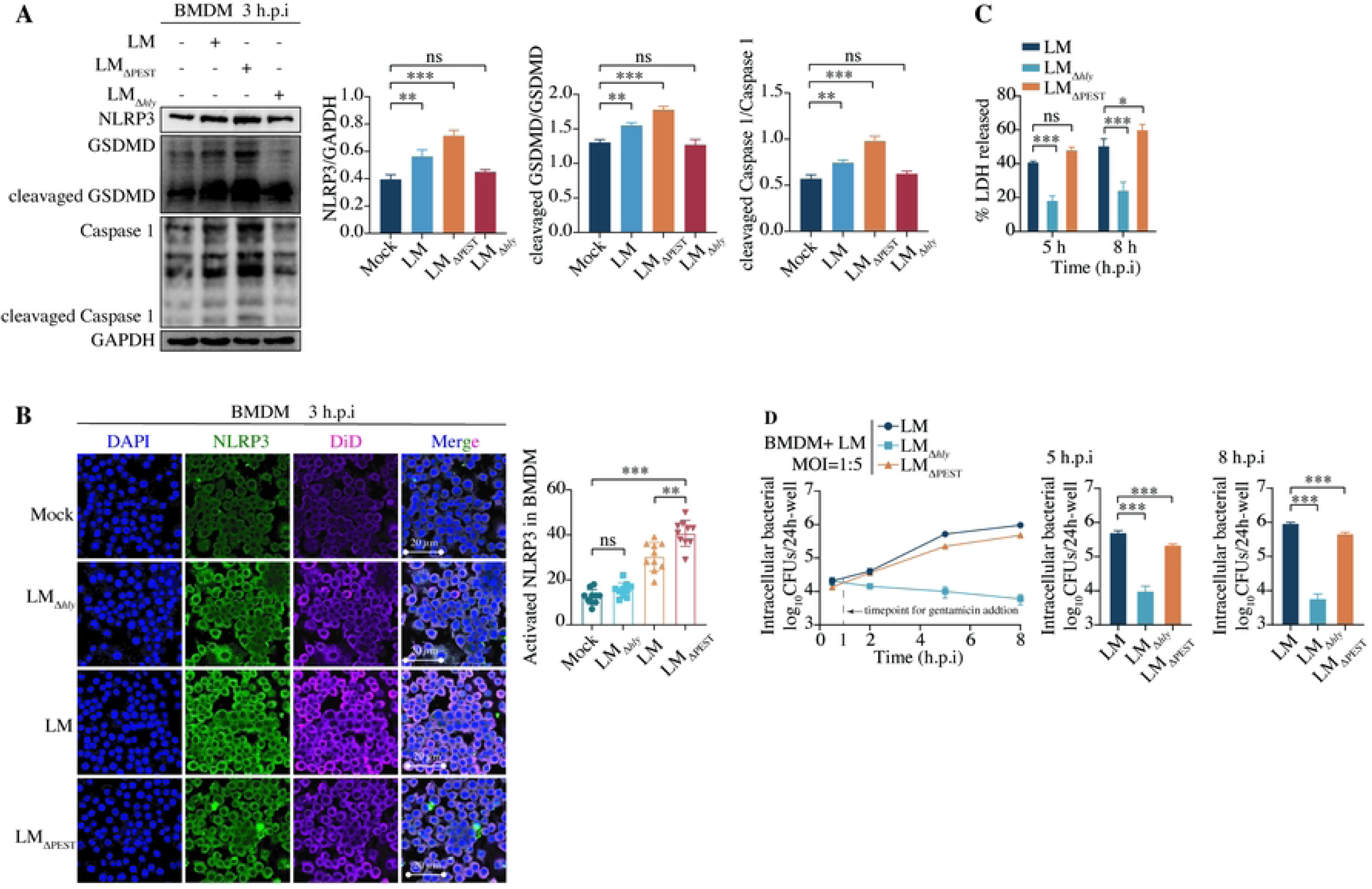
Additional ubiquitination assays support PEST-dependent K48-linked NLRP3 ubiquitination. **(A)** Ubiquitination assay in HEK293T cells co-expressing FLAG-NLRP3, HA-ubiquitin and Myc-LLO, corresponding to the LLO-dependent NLRP3 ubiquitination shown in Fig 3B. FLAG-NLRP3 was immunoprecipitated, and ubiquitination was detected by HA immunoblotting. **(B)** Analysis of ubiquitin linkage specificity using HA-ubiquitin mutants retaining a single lysine residue (K6, K11, K27, K29, K33, K48 or K63) in HEK293T cells expressing FLAG-NLRP3 and Myc-LLO. **(C)** Comparison of normal HA-ubiquitin and K48-only HA-ubiquitin in the presence or absence of Myc-LLO. FLAG-NLRP3 was immunoprecipitated and analysed for ubiquitination. (D) HA-Ub-K48 ubiquitination assay comparing Myc-LLO and Myc-LLO_ΔPEST_ with or without FLAG-NLRP3. Myc-tagged LLO or FLAG-tagged NLRP3 was immunoprecipitated as indicated, followed by immunoblot analysis using an anti-HA antibody. **(E)** Workflow of MYC-LLO affinity purification coupled with mass spectrometry (AP–MS) for identification of LLO-associated proteins. **(F)** KEGG pathway enrichment analysis of LLO-associated proteins. Representative enriched pathways (FDR < 0.05) are shown. **(G)** Biological process classification of proteins identified bv AP-MS. **(H)** Auxiliary prioritisation analysis of degradation- and ubiquitin-system-related candidates, supporting UBA1 as a candidate for further validation. Data are representative of at least three independent experiments.

**Supplementary Fig. 3.**
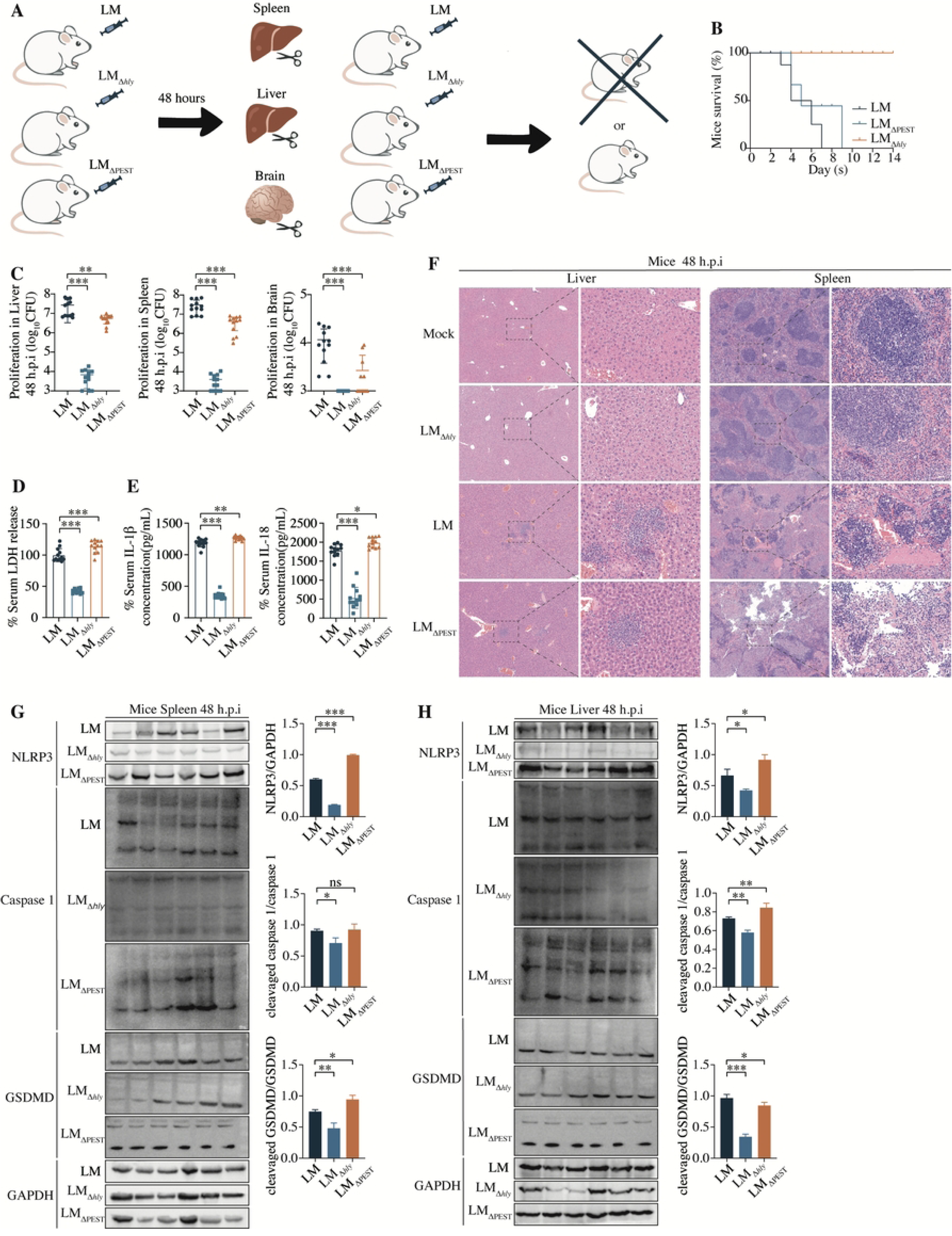
Identification and validation of UBA1 as an LLO-associated ubiquitination factor. **(A)** Effect of increasing UBA1 expression on LLO ubiquitination. HEK293T cells expressing Myc-LLO or Myc-LLO_ΔPEST_ together with HA-ubiquitin and increasing amounts of FLAG-UBA1 (100, 500 and 1,000 ng) were subjected to Myc immunoprecipitation followed by ubiquitin analysis. GAPDH served as a loading control. **(B)** Confocal microscopy analysis of LLO_ΔPEST_ with UBA1 localization in HEK293T cells. Scale bar, 5 μm. **(C)** Interaction analysis between FLAG-NLRP3 and HA-tagged LLO fragments (amino acids 26-193, 194-361 and 362-529) by Co-immunoprecipitation. **(D)** Domain-specific co-immunoprecipitation analysis of UBA1 and NLRP3 domain constructs. **(E-F)** Forward and reciprocal immunoblot-based co-immunoprecipitation analyses of LLO and NLRP3 domain constructs in HEK293T cells. **(G)** Analysis of K48-linked ubiquitination/degradation of NLRP3 domains constructs in the presence of LLO. **(H-I)** Predicted structural models of the LLO-NLRP3 (H) and LLO-UBA1 (I) complexes highlighting putative interaction interfaces. Structural prediction models are used only to suggest possible contact regions and require further validation by mutagenesis and biochemical assays. In A–G, data are from three independent biological replicates.

**Supplementary Fig. 4.**
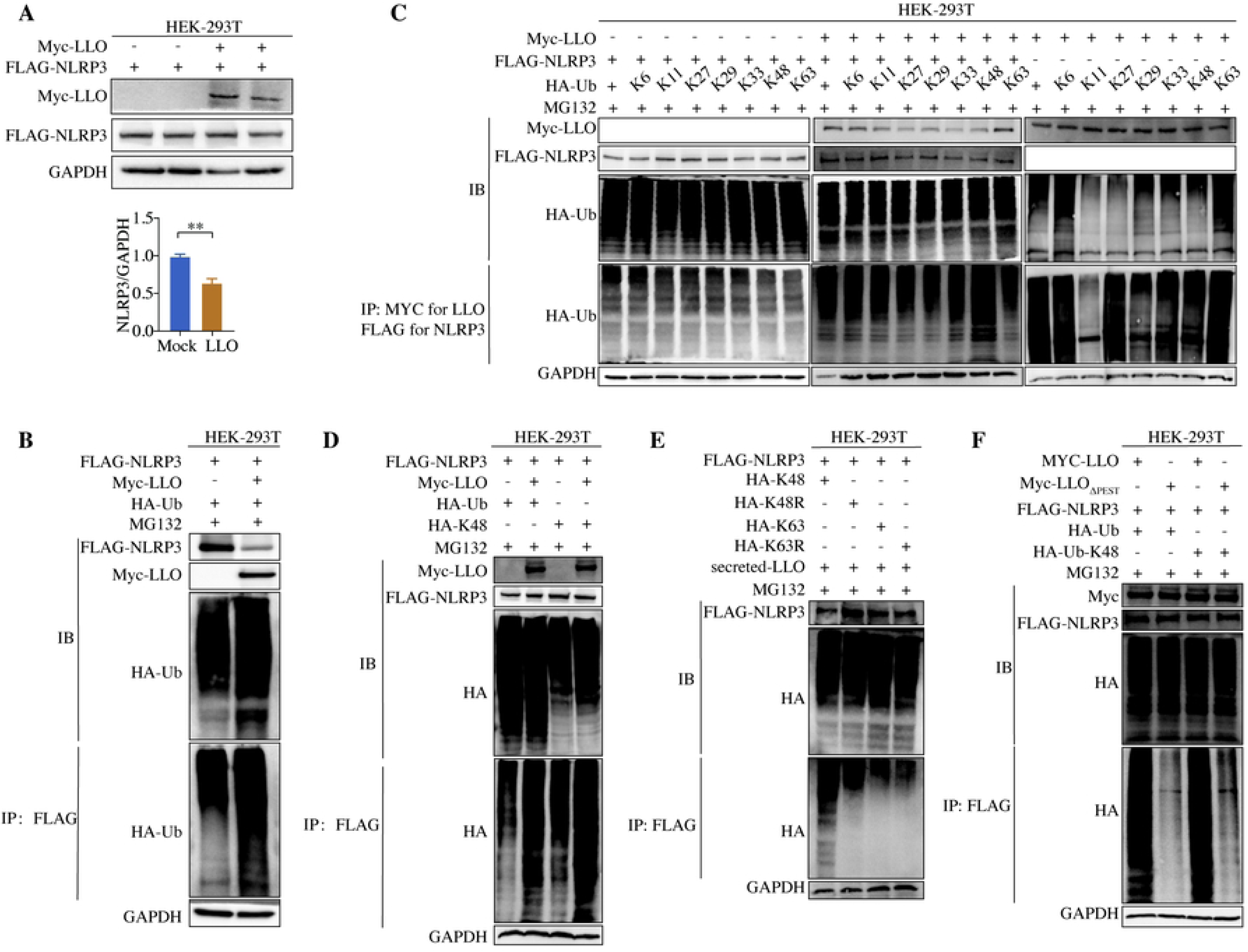
Additional validation of UBA1-dependent regulation of inflammasome activation during *Listeria monocytogenes* infection. **(A)** Gentamicin protection assay measuring intracellular burden of wild-type LM in RAW264.7 cells overexpressing increasing amounts of UBA1 at 5 and 8 h.p.i. **(B)** Immunoblot analysis and densitometric quantification of NLRP3 inflammasome activation in BMDMs overexpressing FLAG-UBA1 following mock, LM_Δ*hly*_ or wild-type LM infection. NLRP3, caspase-1 cleavage and GSDMD cleavage were analysed. **(C)** Immunoblot analysis and densitometric quantification of NLRP3 abundance, caspase-1 cleavage and GSDMD cleavage in BMDMs overexpressing FLAG-UBA1 under mock, wild-type LM or LM_ΔPEST_ infection conditions. **(D-E)** Quantification of IL-1β (D) and IL-18 (E) secretion in BMDMs overexpressing FLAG-UBA1 follwing mock, LM_Δ*hly*_, wild-type LM or LM_ΔPEST_ infection. **(F-G)** Quantitative PCR analysis of IL-1β (F) and IL-18 (G) transcript levels in BMDMs infected with wild-type LM- or LM_Δ*hly*_ following UBA1 overexpression. **(H)** Immunoblot validation of UBA1 knockdown efficiency in BMDMs using independent siRNA sequences at the indicated time points. **(I)** Immunoblot analysis and densitometric quantification of NLRP3 abundance, caspase-1 cleavage and GSDMD cleavage in BMDMs treated with siRNA-UBA1 under mock, LM_Δ*hly*_ or wild-type LM infection conditions. (J) Gentamicin protection assay measuring intracellular bacterial replication in BMDMs expressing full-length NLRP3 or NLRP3 domain-deletion mutants (ΔLRR, ΔPYD or ΔNBD) following LM infection. GAPDH served as a loading control. Data are mean ± s.e.m.; ns, not significant; *p < 0.05, **p < 0.01 and ***p < 0.001. Data are from three independent replicates.

**Supplementary Fig. 5.**
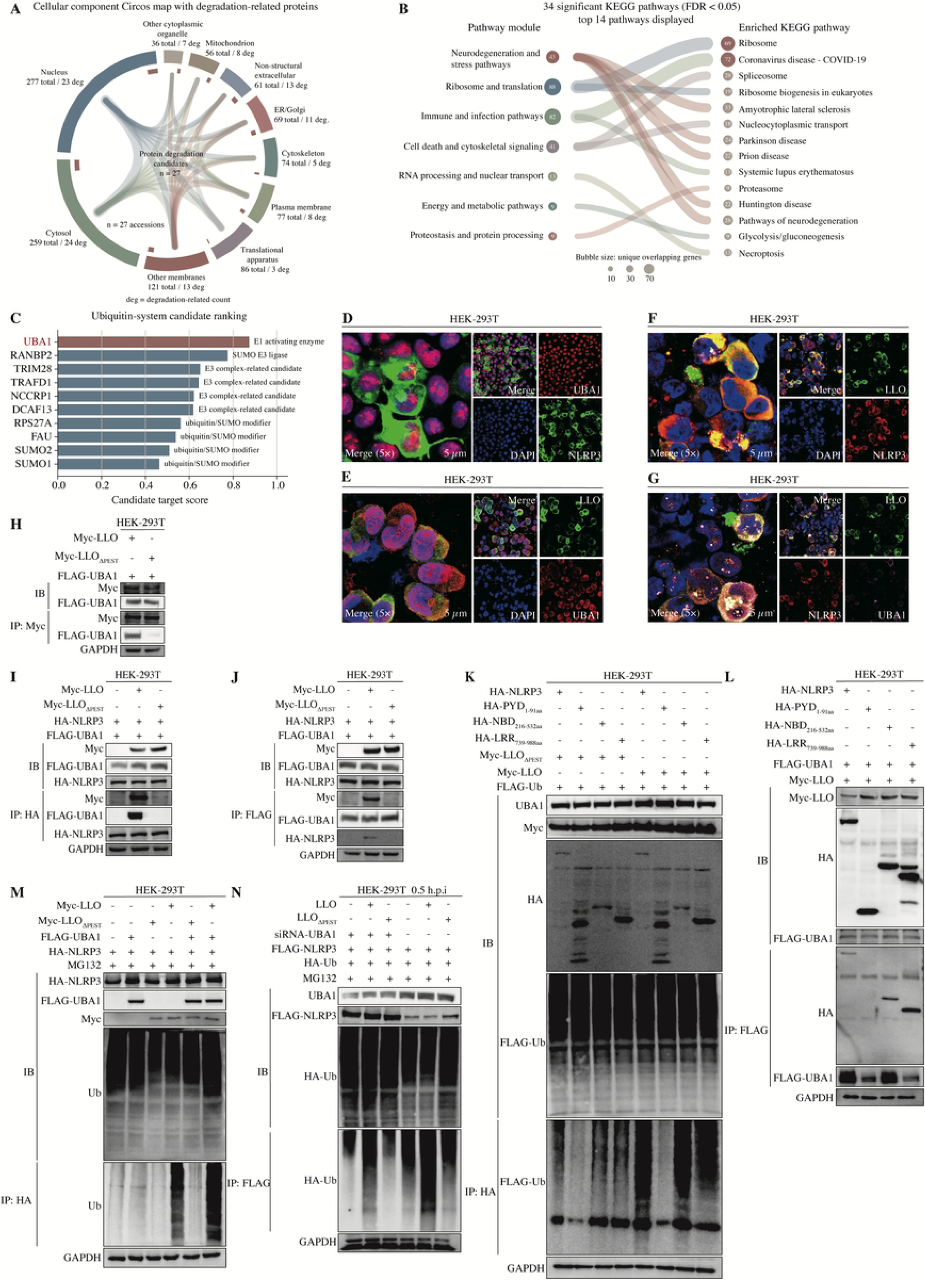
Control analyses for *in vivo* UBA1 knockdown and LLO dependence. **(A-B)** Immunoblot analysis of UBA1, NLRP3, caspase-1, cleavaged caspase-1, GSDMD and cleavaged GSDMD in the spleen (A) and liver (B) from siRNA-UBA1-treated or control mice at 48 h.p.i. GAPDH served as the loading control. **(C)** Representative haematoxylin and eosin staining of the liver and spleen from mock-treated mice, LM_Δ*hly*_ -infected mice, siRNA-UBA1-treated mice and siRNA-UBA1-treated mice infected with LM_Δ*hly*_ at 48 h.p.i. Scale bars, 0.1 mm for low-magnification images and 0.02 mm for high-magnification images. Data are representative of at least three independent experiments.

## Notes

### Competing Interest Statement

The authors have declared no competing interest.

